# Ketogenic diet alters microglial morphology and changes the hippocampal lipidomic profile distinctively in stress susceptible *versus* resistant male mice upon repeated social defeat

**DOI:** 10.1101/2023.08.28.555135

**Authors:** Fernando González Ibáñez, Torin Halvorson, Kaushik Sharma, Chloe McKee, Micaël Carrier, Katherine Picard, Nathalie Vernoux, Kanchan Bisht, Jessica Deslauriers, Maciej Lalowski, Marie-Ève Tremblay

## Abstract

Psychological stress confers an increased risk for several diseases including psychiatric conditions. The susceptibility to psychological stress is modulated by various factors, many of them being modifiable lifestyle choices. The ketogenic diet (KD) has emerged as a dietary regime that offers positive outcomes on mood and health status. Psychological stress and elevated inflammation are common features of neuropsychiatric disorders such as certain types of major depressive disorder. KD has been attributed anti-inflammatory properties that could underlie its beneficial consequences on the brain and behavior. Microglia are the main drivers of inflammation in the central nervous system. They are known to respond to both dietary changes and psychological stress, notably by modifying their production of cytokines and relationships among the brain parenchyma. To assess the interactions between KD and the stress response, including effects on microglia, we examined adult male mice on control diet (CD) *versus* KD that underwent 10 days of repeated social defeat (RSD) or remained non-stressed (controls; CTRLs). Through a social interaction test, stressed mice were classified as susceptible (SUS) or resistant (RES) to RSD. The mouse population fed a KD tended to have a higher proportion of individuals classified as RES following RSD. Microglial morphology and ultrastructure were then analyzed in the ventral hippocampus CA1, a brain region known to present structural alterations as a response to psychological stress. Distinct changes in microglial soma and arborization linked to the KD, SUS and RES phenotypes were revealed. Ultrastructural analysis by electron microscopy showed a clear reduction of cellular stress markers in microglia from KD fed animals. Furthermore, ultrastructural analysis showed that microglial contacts with synaptic elements were reduced in the SUS compared to the RES and CTRL groups. Hippocampal lipidomic analyses lastly identified a distinct lipid profile in SUS animals compared to CTRLs. These key differences, combined with the distinct microglial responses to diet and stress, indicate that unique metabolic changes may underlie the stress susceptibility phenotypes. Altogether, our results reveal novel mechanisms by which a KD might improve the resistance to psychological stress.

**Highlights:**

- Ketogenic diet tends to promote resistance to psychological stress
- Hippocampal microglia show morphological adaptations to stress and diet
- Microglia of stress-susceptible mice make less synaptic contacts
- Microglia of ketogenic diet-fed mice show less signs of cellular stress
- Lipids are differentially regulated in the hippocampi of susceptible mice

## 1. Introduction

In recent years, dietary interventions have garnered increased scientific and clinical interest for the treatment of psychiatric disorders, including major depressive disorder (MDD) ^1–7,7–9^. In particular, the ketogenic diet (KD), already established as a treatment for epilepsy ^1,10^, has gained attention for psychiatric disorders, as well as for promoting optimal cognition in healthy individuals ^5,8,11–13^. A standard KD consists of up to 80% of lipids, with adequate protein and minimal carbohydrate contents ^5,6,8^. Contrary to a typical carbohydrate-rich diet, in which glucose is the primary energy source for most tissues, including the brain ^1,14^, a KD regimen forces the body into ketosis, a metabolic state in which lipids are converted to ketone bodies, such as β-hydroxybutyrate (BHB) and acetoacetate, in the absence or reduced presence of glucose ^1,8^. Energy is derived from the catabolism of fatty acids and ketone bodies, with the brain relying heavily on ketone bodies ^1,8,15^. A strong anti-oxidative effect has been proposed to underlie the beneficial effects of KD in epilepsy ^16^. Furthermore, the metabolic changes induced by a KD regimen may also exert positive physiological and cognitive outcomes under normal homeostatic conditions ^12^. Thus, a KD may represent a potential approach to prevent the emergence of psychiatric disorders in healthy individuals, notably considering the preclinical evidence suggesting that a KD may promote resilience to stress-induced depression ^7^.

Recent preclinical and clinical reports have revealed the potential of a KD to exert strong antidepressant effects ^4,7^. In a model of repeated social defeat (RSD), a KD ameliorated depressive-like behavior in stress-susceptible mice, as well as improved the social interaction ratios, sucrose preference and performance in the tail suspension and forced swim tests ^7^. A prior study on adult male mice demonstrated significant reductions in anxious and depressive-like behaviors following two weeks of KD ^4^. Evidence from humans is still limited, but improvements in mood, cognitive function, and anxious behavior were noted in young patients undergoing KD therapy for epilepsy ^17^. Compromised hippocampal function is thought to underlie the classical symptoms of depression such as impaired concentration and declarative memory, and affective changes accompanied by anxiety ^18,19^. Significant reductions in the volume of the ventral hippocampus, involved in anxiety ^20^, were reported in rodent models of environmental challenges including mice subjected to psychological stress ^21,22^. Similarly, reduction in hippocampal volume has been reported in human patients with MDD ^23,24^. Synaptic loss was identified as a driver of hippocampal volume reduction in mouse models of chronic stress ^25,26^. This is supported by observations in patients with MDD where volume reduction ^27^ and synaptic loss ^28^ in the hippocampus were associated with MDD severity.

Acute and chronic stress, but also cumulative stress exposure, are major risk factors for MDD ^29,30^. Recent work has demonstrated that psychological stress and depression are strongly linked to inflammation, which is thought to play a causal role at least in a subset of depression cases ^31–36^. Notably, RSD stress in mice triggers central and peripheral inflammatory processes, inducing similar depressive-like behaviors ^7^, which further supports a causal link between inflammation and depression. Elevated basal levels of inflammatory markers were also linked to depressive symptoms in humans ^37–41^. Conversely, anti-inflammatory interventions, such as dietary regimens, may benefit patients with MDD who display chronic inflammation ^42–44^. Among the underlying mechanisms, microglia, the resident innate immune cells of the central nervous system, are key mediators of brain inflammation ^39,45–48^. Emerging evidence suggests that microglia play an important role in the pathophysiology of stress and depression, particularly as mediators of pathological inflammation, but also vascular, neuronal, and synaptic remodeling ^49–51^. Morphological and functional alterations of microglia were described notably among the hippocampus in mouse models of RSD ^7,52^. Similar observations were made in patients with MDD ^53^. The microglial alterations observed in rodent models of psychological stress include accelerated cellular aging, as well as senescence and metabolic dysregulation, associated with energetic deficits ^46,54^. Stress compromises the physiological role of microglia in maintaining homeostasis, resulting in increased basal levels of inflammatory cytokines in the brain, considered to be detrimental to microglia-neuron interactions and cognitive function ^46^. However, microglial release of anti-inflammatory cytokines and neurotrophic factors, which promote adult hippocampal neurogenesis ^50,51,55^, among other beneficial functions, may contribute to counteracting some of these changes.

The evidence linking inflammation with chronic psychological stress and depression has prompted an interest in developing therapeutic interventions acting on inflammation, to treat depressive disorders and favor optimal cognitive health ^50,56,57^. Among the proposed strategies, a KD regimen was shown to exert anti-inflammatory effects, by reducing the peripheral and central levels of pro-inflammatory cytokines notably in response to RSD in adult male rats and mice ^7,58^. Furthermore, the major ketone body BHB is a known inhibitor of the nucleotide-binding and oligomerization domain-like receptors pyrin domain-containing protein 3 (NLRP3) inflammasome, which acts as an important mediator of inflammation in innate immune cells that include microglia ^59,60^. Microglia are emerging as a critical mediator of several beneficial effects of a KD, such as reducing brain inflammation and improving depressive behaviors in humans ^50^, while changes in microglia immune-metabolic pathways have been highlighted as a central mechanism underlying MDD ^61^. A KD is thus hypothesized to attenuate stress-related cellular aging and inflammation through the normalization of microglial metabolism and functions.

To provide further insight into the outcomes of KD (*versus* a control diet; CD) on stress resilience, inflammation, and microglia, we utilized a RSD paradigm in adult male mice. The study was performed on the ventral hippocampus CA1 *stratum radiatum* considering its key role in the plasticity impairment observed upon chronic social stress and in MDD ^20,22–24,62–64^. We first assessed the prevalence of susceptible *versus* resistant mice under CD and KD following the RSD. We then compared blood levels of anti- and pro-inflammatory mediators with the diets at steady-state and after psychosocial stress. Furthermore, we characterized changes in hippocampal microglial density, morphology, and ultrastructure (including organelles and relationships with parenchymal elements such as synapses), as well as analyzed the hippocampal lipidomic profile under KD *versus* CD at steady-state and in the context of psychosocial stress.

## 2. Methods

### 2.1 Animals

All animal experiments were performed under approval of the institutional animal ethics committees, in conformity with the Canadian Council on Animal Care guidelines. Male mice were used considering that RSD relies on inter-male interactions ^65,66^. C57BL/6J mice (7–8 weeks old) were acquired from The Jackson Laboratories and CD1 retired breeder mice (4–6 months old) from Charles River (St. Constant, QC, Canada). A total of 82 animals were used in this study. The animals were housed under a 12 h light-dark cycle at 22–25 °C with *ad libitum access* to food and water.

### 2.2 Ketogenic diet

Starting 4 weeks prior to the RSD paradigm, the experimental C57BL/6J mice gradually transitioned to KD or remained on CD over 1 week (Fig. 1A). The KD (high fat and low carbohydrate content) had a composition of 8.6% protein, 75.1% fat, 4.8% fiber, 3.2% carbohydrate, caloric profile: protein 0.34 kcal/g, fat 6.76 kcal/g and carbohydrate 0.13 kcal/g [Ketogenic Diet AIN-76A-Modified, High Fat, F3666 Bio-Serve]. The CD was composed of 24% protein, 18% fat, 58% carbohydrate, caloric profile of protein 0.744 kcal/g, fat 0.558 kcal/g and 1.798 kcal/g [Teklad Global 18% protein, 2018S, ENVIGO]. The mice were weighted daily.

**Figure 1:**
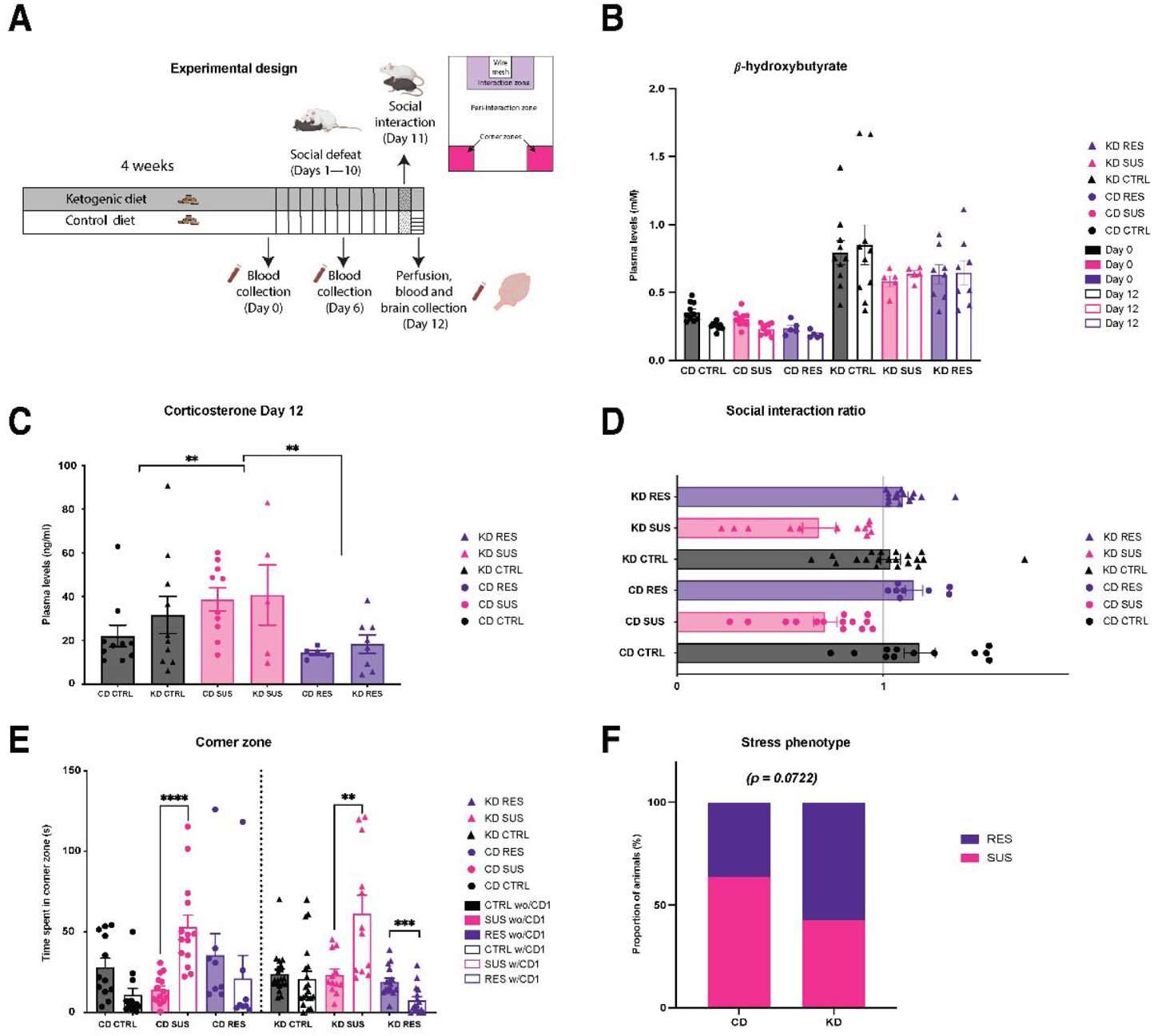
Ketogenic diet tends to increase stress resistance in a mouse model of repeated social defeat. Experimental timeline, mice were fed 4 weeks of ketogenic diet (KD) in KD group, or control diet (CD) in CD group. Mice were exposed to ten days of repeated social defeat (RSD), first day of social defeat is considered Day 1. Social interaction (SI) test was performed on Day 11. Blood draws were performed on Days 0, 6 and 12. Perfusion and brain collection were performed on Day 12 **(A)**. Four weeks of KD increased blood levels of β-hydroxybutyrate (BHB) throughout RSD until brain collection **(B)**. Susceptible (SUS) mice on CD and KD had increased blood levels of CORT corticosterone compared to control (CTRL) and resistant (RES) mice **(C)**. SUS mice on CD and KD showed increased levels of social avoidance in a SI test **(D)**. SUS mice on CD and KD spent more time in corner zone (CZ) once the CD1 mouse entered the arena **(E)**. RES mice on KD spent less time in CZ once the CD1 entered the arena. Mice on KD *versus* CD showed an increased proportion of mice classified as RES **(F)**. n = 5–10 mice/group for the BHB and corticosterone tests; n = 8–20 mice/group for the behavioral tests. Data are expressed as mean ± standard error of the mean. Statistical significance was assessed by 2-way ANOVA followed by Tukey *post-hoc* analysis, where **p < 0.01 and ****p < 0.0001. For corticosterone analysis, statistical significance was assessed by 1-way ANOVA, where **p < 0.01. CD: control diet; KD: ketogenic diet; CTRL: control; SUS: susceptible; RES: resistant. Created with the help of BioRender.

### 2.3 Blood samples

Blood samples were collected from all the experimental C57BL/6J mice through the mandibular vein, without anesthesia, on Day 0, Day 6 of the social defeat and on Day 12 of the social interaction (SI) test (Fig. 1A). Blood was collected in heparinized tubes and centrifuged at 3600 revolutions per min for 10’ at 4°C to collect plasma.

#### 2.3.1 Corticosterone measurement

Plasma corticosterone (CORT) levels were determined using blood samples collected on Day 12 using a commercial CORT ELISA kit (item No. 501320, Cayman Chemical, Ann Arbor, MI, USA), according to the manufacturer’s instructions. Plates were read at 405 nm with a microplate reader (iMarkTM, Biorad, Hercules, CA, USA). Sample concentrations were determined using a standard curve (logarithmic scale), followed by a four-parameter logistic fit analysis (Henry et al. 2018).

#### 2.3.2 b-hydroxybutyrate measurement

Plasma BHB levels were determined to confirm the increase of circulating ketone bodies in the KD fed animals, using a β-hydroxybutyrate or 3-hydroxybutyric acid Colorimetric Kit Essay (item No. 700190Cayman, Chemical, Ann Arbor, MI, USA), according to the manufacturer’s instructions. Absorbance was read at 445–455 nm using a plate reader.

#### 2.3.3 Cytokine measurement

Plasma levels of 31 cytokines were measured with the Discovery Assay® by Eve technologies (Mouse Cytokine/Chemokine 31-Plex Discovery Assay® Array (MD31); Calgary, Canada). These cytokines were as follows: eotaxin, granulocyte colony stimulating factor (G-CSF), granulocyte macrophage colony stimulating factor (GM-CSF), interferon (IFN)gamma, interleukin (IL)-1α, IL-1β, IL-2, IL-3, IL-4, IL-5, IL-6, IL-7, IL-9, IL-10, IL-12 (p40), IL-12 (p70), IL-13, IL-15, IL-17A, C-X-C motif chemokine ligand 10 (IP-10), keratinocytes-derived chemokine (KC), leukemia inhibitory factor (LIF), lipopolysaccharide-induced CXC chemokine (LIX), monocyte chemoattractant protein-1 (MCP-1), macrophage colony stimulating factor (M-CSF), monokine induced by interferon-γ (MIG), macrophage inflammatory protein 1-alpha (MIP-1α), macrophage inflammatory protein 1β (MIP-1beta), macrophage inflammatory protein 2 (MIP-2), C–C chemokine ligand 5 (RANTES), TNF-α, and vascular endothelial growth factor (VEGF).

### 2.4 Social defeat

#### 2.4.1 Social defeat paradigm

CD1 retired breeder mice were selected for their level of aggressiveness in presence of naïve C57BL/6J mice as previously published by Golden *et al*., 2015 and Henry *et al.,* 2018. Experimental C57BL/6J mice were randomly assigned to the RSD or non-stressed control (CTRL) group. Experimental mice (intruder) were subjected to 10 days of consecutive RSD (Day 1 is the first day of RSD). The social defeat arena is a cage with two equal sections divided by a physical barrier with holes that allow for visual, olfactory, and auditory interactions between the sections. An aggressor CD1 mouse (resident mouse) is first placed on one side of the barrier, then an intruder mouse is placed for 5 min with the resident mouse. After 5 min, the intruder mouse is moved to the other side of the barrier for the next 24 h until the intruder mouse is exposed to a new resident mouse (and the resident mouse to a new intruder) ^65,66^. Each intruder was randomly exposed to the same group of aggressor mice in a different order. The health status of the intruder mice was monitored closely. Injuries are not required for this protocol. In order to control for unnecessary physical lesions, the social defeat was interrupted after 10 physical attacks within a session or when attacks lasted more than 3 seconds. After every social defeat, the mice were examined by animal care technicians, and their wounds were treated. In our study, only one mouse was heavily wounded, then euthanized and excluded from the experiment according to the veterinarian guidelines.

For non-stressed controls, the mice were paired with partners, and housed in the same type of cage changed daily in the same room as RSD animals. All groups, stressed and non-stressed on both diets, underwent the SI test. SI test was performed on Day 11 and animal euthanasia on Day 12. The animal movement was tracked in the arena, before and after the introduction of a novel CD1 aggressor. Using the time mice spent in each zone, a SI ratio was determined as described below.

#### 2.4.2 Social interaction test

One day after the final defeat (Day 11), experimental animals were placed alone inside an open field arena (42 cm x 42 cm x 42 cm) for 150 s. The social interaction arena was divided into corner zones (CZ), an interaction and peri-interaction zones (Fig. 1A). After 150 s, an aggressor mouse not previously used in the RSD paradigm was placed inside a wire mesh in the arena, contained in the interaction zone (IZ). The aggressor mouse remained there for 150 s. Tracking videos were captured and analyzed using ANY-maze (Stoelting Co, Wood Dale, IL, USA). The test was performed by an observer blind to the experimental conditions. The SI ratio was calculated by dividing the time spent in the interaction zone and peri-interaction zone in the presence *versus* absence of the CD1 mouse. Susceptible mice (SUS) tended to freeze, avoiding interaction with the aggressor mouse, and spending more time in the CZs, thus showing social avoidance. Resistant mice (RES) still interacted with the aggressor mice, staying in the interaction and peri-interaction zones, adapted from ^65,66^ (Fig. 1A). To classify mice either as RES or SUS, a theoretical cut-off of 1 was used for the SI ratio. Mice with a SI ≥1 were considered as RES and mice with SI ratio < 1 were considered as SUS ^65,66^. CTRL mice showing a SI <1 were excluded, following ^33,66–68^. Excluded CTRL mice with a SI < 1: CD = 2 mice; KD = 9 mice.

#### 2.4.2 Euthanasia and tissue preparation

On Day 12 (i.e., 1 day after the SI test to exclude acute effects of the social interaction), mice were anesthetized with ketamine and xylazine (80 and 10 mg/kg, respectively, by intraperitoneal injection), and euthanized (Fig. 1A). For imaging techniques, mice were perfused with phosphate-buffered saline (PBS; 50 mM pH 7.4) followed by a mixture of 4% paraformaldehyde (PFA) and 0.2% glutaraldehyde in phosphate buffer (PB; 100 mM, pH 7.4). Using a vibratome (Leica VT1000S), 50 µm thick coronal sections were obtained and stored in cryoprotectant at −20 °C until use. For lipidomics, the mice were perfused with PBS, the hippocampus was collected and flash-frozen on dry ice and stored at −80 °C. To allocate mice to different experiments (e.g., microscopy *versus* omics), this was planned beforehand given that different experiments required different perfusion methods. The animals were randomly assigned to an experiment. The only experiment where mice were selected was for lipidomics, samples from the 3 mice presenting the strongest phenotype, based on their SI ratio, were sent for analysis. There was no calculation of the average, all SUS and RES animals were treated equally, except for lipidomics, as mentioned previously.

### 2.5 Density and morphology

#### 2.5.1 Fluorescence staining

Fluorescence staining was performed as described in Gonzalez-Ibanez *et al*., 2019. Utilizing 3–5 animals per experiment group, 2–5 sections containing the ventral hippocampus corresponding to Bregma levels −2.88 to −3.38 mm were elected based on the stereotaxic atlas of Paxinos and Franklin (4^th^ edition). Sections were washed with PBS five times for 5 min and incubated with 10 mM citrate buffer at 70 °C. After reaching room temperature (RT), they were washed with PBS five times for 5 min and incubated in 0.1% NaBH_4_ for 30 min followed by 5 times for 5 min washes with PBS. Sections were then incubated in a blocking buffer (BB, 0.5% gelatin, 5% normal goat serum, 5% normal donkey serum and 0.01% Triton X-100) for 1 h. Subsequently, sections were incubated overnight at 4 °C with BB containing a primary antibody cocktail (1:150 mouse anti-ionized calcium-binding adapter molecule 1 (Iba1; EMD-Millipore cat# MABN92+), 1:300 rabbit anti-transmembrane protein 119 (TMEM119; Abcam cat# ab209064)). The next day, the sections were washed with PBS containing Triton X-100 (PBST, 0.01% Triton X-100 in PBS) five times for 5 min, followed by an incubation in BB with a secondary antibody cocktail. For the morphology analysis, we used: 1:300 donkey anti-mouse Alexa555 (Invitrogen-ThermoFisher cat# A-31570) and 1:300 goat anti-rabbit Alexa647 (Invitrogen-ThermoFisher cat# A-21245). For the density and distribution analysis: 1:300 donkey anti-mouse Alexa555 (Invitrogen-ThermoFisher cat# A-31570), 1:300 goat anti-rabbit Alexa647 (Invitrogen-ThermoFisher cat# A-21245) or 1:300 donkey anti-mouse Alexa488 (Invitrogen-ThermoFisher cat# A-21202) and 1:300 goat anti-rabbit Alexa568 (Invitrogen-ThermoFisher cat# A-11011) for 90 min. Sections were then washed five times for 5 min with PBST, incubated with DAPI 1:20,000 for 5 min and washed with PB three times for 5 min. The sections were mounted, dried overnight and coverslipped with mounting medium Fluoromount G (cat# 0100-01, Southern-Biotech, Birmingham, AB, USA).

#### 2.5.2 Microglial density and distribution

Density and distribution analyses were performed in the ventral hippocampus CA1 *stratum radiatum* where stress-driven microglia-synapse interactions were previously studied ^69^. In each of 3 animals per experimental group, 2–3 sections containing the ventral hippocampus CA1 *stratum radiatum* were used to build a mosaic at 20x. Images were acquired with an Axio Imager M2 epifluorescence microscope equipped with an AxioCam MRm camera using the Zen Pro 2012 software (Zeiss, Oberkochen, Germany). With ImageJ, the freehand tool was used to delimit the region of interest (ROI) which was then measured in mm^2^. Using DAPI as a confirmation of cellular identity, all Iba1+/TMEM119+ cells were considered as microglia and all Iba1+/TMEM119-cells were considered as peripheral infiltrating macrophages ^70,71^. Using the paintbrush tool, all cell bodies were registered and quantified. A total of 200–300 cells per animal was included in the analysis. To assess density, the total number of microglia was divided by the measured area. Using ImageJ’s nearest neighbor distance (NND) plug in, the distance (µm) of each microglia to its closest neighbor was obtained. The average NND of all microglia per ROI was calculated. Spacing index was calculated as the square average NND multiplied by the density (arbitrary units). This value was averaged across mosaics to determine the value per animal as in Tremblay *et al.* 2012 and Ibáñez *et al.* 2019.

#### 2.5.3 Microglial morphology

For morphology analysis, 3–5 animals per experimental group were utilized. In each animal, 14– 19 microglia (Iba1+/Tmem119+ cells) from the ventral hippocampus CA1 *stratum radiatum* were randomly selected and analyzed, resulting in 51–88 cells/experimental group, a sample size which was considered sufficient to obtain statistical power based on the G*Power software V3.1 (effect size of 0.231 and power of 0.95 estimated to a total of 378 individual cells). Cells were imaged at 40x with a Z-interval of 0.33 µm using a Quorum WaveFX spinning disc confocal microscope (Quorum Technologies, Guelph, ON, Canada) equipped with an ORCA-R2 camera (512 x 512 pixels; Hamamatsu Photonics, Hamamatsu, Japan). A Z-project projection maximum intensity image was generated using the ImageJ Z-stack tool. Morphological analysis was done using the Iba1 channel in ImageJ. Using the ImageJ freehand tool, the soma of each cell was traced and measured in µm^2^. With the ImageJ polygon tool, the arborization area was traced by selecting the tips of microglial processes and then measured in µm^2^. The morphological index was performed by dividing the soma area by the arborization area, as performed in Tremblay *et al.* 2012 and González Ibáñez *et al.* 2019.

### 2.6 Ultrastructural analysis

#### 2.6.1 Microglial ultrastructural analysis

In each of 3 animals per group, 3 sections containing the ventral hippocampus CA1 *stratum radiatum* were selected. Sections were washed with PBS five times for 5 min. Samples were post-fixed with 3.5% acrolein in PB for 2 h. Sections were washed with PBS five times for 5 min and incubated in 0.3% NaBH_4_ for 30 min followed by five times for 5 min washes with PBS. Sections were then incubated in BB (10% fetal bovine serum, 3% bovine serum albumin, 0.01% Triton-X) for 1 h. Subsequently, the sections were incubated overnight at 4 °C in BB with primary antibody cocktail ([1:1000] rabbit anti-Iba1 polyclonal primary antibody (FUJIFILM, Wako Chemical, Osaka, Japan, cat#019-19741). After reaching RT, the sections were washed with Tris-buffered saline (TBS, 50 mM, pH 7.4) five times for 5 min and incubated in BB containing biotinylated goat anti-rabbit polyclonal secondary antibody ([1:200] Jackson ImmunoResearch, West Grove, PA, USA, cat# 111-066-046) in TBS for 1.5 h. Sections were next incubated with avidin-biotin complex solution (Vector Laboratories, Burlingame, CA, USA, cat# PK-6100,) [1:100] in TBS; for 1 h at RT. The staining was revealed in 0.05% diaminobenzidine (DAB; Millipore Sigma cat# D5905-50TAB,) with 0.015% H_2_O_2_ in Tris-buffer (TB, pH 8.0) for 4.5 min at RT. Samples were post-fixed in osmium-thiocarbohydrazide-osmium to enhance contrast for scanning electron microscopy (SEM). Sections were incubated in a 1:1 solution of 4% aqueous osmium tetroxide (Electron Microscopy Sciences (EMS), Hatfield, PA, USA cat#19170) and 3% potassium ferrocyanide (Bio-Shop, Burlington, ON, Canada, cat# PFC232.250) in double distilled (dd)H_2_O for 1 h. Sections were washed with ddH_2_O three times 5 min and incubated in 1% thiocarbohydrazide (EMS, cat# 2231-57-4) diluted in ddH_2_O for 20 min. After washing the sections three times for 5 min, they were incubated for 30 min in 2% osmium tetroxide diluted in ddH_2_O and then dehydrated in ascending concentrations of ethanol (two times in 35%, one time in 50%, 70%, 80%, 90%, three times 100%) followed by three incubations of 5 min in propylene oxide. After dehydration, the sections were flat-embedded in Durcupan ACM resin (Millipore Sigma, cat# 44611-44614). In brief, the sections infiltrated the resin at RT overnight. They were carefully placed on a fine layer of resin between 2 sheets of ACLAR® embedding films (EMS, cat# 50425-25) for polymerization at 55 °C for 72 h. After polymerization, a section containing the region of interest was excised and glued to a Durcupan resin block for ultrathin sectioning (Ultracut UC7 ultramicrotome, Leica Biosystems). Ultrathin sections, of ~75 nm thickness, were collected on a silicon nitride chip and placed on specimen mounts for SEM. In each animal, 10–14 randomly selected microglial cell bodies located in the CA1 *stratum radiatum* were imaged, resulting in a total of 33–38 cells per condition, a sample size which was considered sufficient to obtain statistical power based on the G*Power software V3.1 (effect size of 0.313 and power of 0.95 estimated to 210 individual cells). The cells were imaged at 5 nm of resolution using a Crossbeam 540 field emission SEM with a Gemini column (Zeiss). The quantitative analysis was performed blind to the experimental conditions using QuPath Software.

#### 2.6.2 Ultrastructural identification

Microglial cell bodies were identified by their dark irregular cytoplasm, heterogeneous chromatin pattern, distinctive long stretches of endoplasmic reticulum (ER) and lipidic inclusions (i.e., lipofuscin, lipid bodies or droplets, lysosomes), as well as frequent contacts with axon terminals ^72,73^. Contacts with blood vessels, astrocytic cell bodies and neuronal cell bodies were quantified. Neurons were identified by their pale nuclei, pale cytoplasm, common presence of a nucleolus and their round shape with a frequent apical dendrite or axon projecting from the cell body. Astrocytes were identified by their pale nuclei, a fine rim of heterochromatin lining the nuclear membrane, acute angles, and frequent intermediate filaments, among other features. Blood vessels (BV) were identified by their lumen, endothelial cells, and surrounding basal membrane. Contacts to blood vessels were considered when microglial cell body was directly touching the basal membrane of the BV or in proximity to the basement membrane ^72,73^. Microglial contacts with other neuronal structures, particularly pre-synaptic axon terminals and post-synaptic dendritic spines, were quantified. Pre-synaptic axon terminals were identified by their synaptic vesicles, with a minimum of 5 vesicles required for recognition. Post-synaptic dendritic spines were identified by their post synaptic density and apposition with a pre-synaptic axon terminal ^72,73^. Contacts to synaptic clefts were considered when microglial cell bodies directly juxtaposed both excitatory synapse-forming elements ^74,75^. Microglial-synaptic contacts were classified as axon terminals, dendritic spines, or synaptic clefts.

Microglial mitochondria, ER, Golgi apparatus, lysosomes, lipofuscin granules, nuclear membrane alterations, nuclear pores and autophagosomes were quantified and their health status was assessed ^73^. Mitochondria longer than 1 µm were considered elongated ^75^. Mitochondria containing electron-lucent circular hollow membrane rings were categorized as “holy” mitochondria ^75^. Swollen mitochondria with abnormal cristae structure were considered dystrophic mitochondria. Mitochondria with clear appearance and small fractured cristae were defined as white mitochondria ^73^. Total dystrophic mitochondria count was obtained by adding dystrophic mitochondria, white mitochondria and holy mitochondria. Mitochondria without any of these alterations were considered as standard mitochondria. A percentage of dystrophic mitochondria was calculated based on total mitochondria count. Dilation of the ER was noted when the cisternae had an electron-lucent appearance and the intracisternal distance was 100 nm or higher ^76,77^. The presence of inclusions refers to electron-dense material within the intracisternal space. ER without signs of dilation or inclusions was considered standard ER. The total dystrophic ER number was calculated by the sum of total ER with dilation and inclusions. A total ER count was calculated by the sum of standard ER and total dystrophic ER. Percentage of dystrophic ER was calculated based on total ER count. Dilation of the Golgi apparatus was noted when the cisternae had an electron-lucent appearance and the intracisternal distance was 100 nm or more ^76,77^. Inclusions refer to the accumulation of electron-dense material within the intracisternal space. Golgi apparatus without signs of dilation or inclusions was considered standard Golgi apparatus. Total dystrophic Golgi apparatus was calculated by the sum of total Golgi apparatus with dilation and inclusions. A total Golgi apparatus count was calculated by the sum of standard Golgi apparatus and total dystrophic Golgi apparatus. Percentage of dystrophic Golgi apparatus was calculated based on the total Golgi apparatus count. Lipid inclusions were identified by their round shape and electron-dense color and smooth texture. Lipofuscin granules were identified by their round or oval shape, and granular appearance with thread-like structures resembling a fingerprint-like pattern ^78^. Phagosomes were quantified, discriminating between empty phagosomes and phagosomes with content ^76^. The presence of content was defined as electron-dense material contained in the phagosome. Total phagosomes were calculated by adding empty phagosomes and phagosomes with content. A percentage of phagosomes with content was calculated based on the total number of phagosomes. Primary lysosomes were recognized by their dense homogeneous salt and pepper texture, round shape, and single membrane enclosure ^75,79^. Secondary lysosomes were identified by their association with endosomes, small lipid droplets and inhomogeneous texture. Tertiary lysosomes were identified by their association to lipofuscin granules and lipid droplets ^75,78,79^. A total lysosomal count was calculated by adding primary, secondary and tertiary lysosomes. Nuclear pores were identified as an interruption of the nuclear membrane, when the outer and inner nuclear membranes were joined. Nuclear indentations were defined as an invagination of the nuclear membrane. Nuclear alterations consisted in an alteration of the nuclear integrity or presence of inclusions within the nuclear membrane ^73^. All data was registered per cell and averaged by experimental condition.

### 2.7 Lipidomic analysis

#### 2.7.1 Liquid chromatography/mass spectrometry (LC/MS)

Whole hippocampi were collected from 4 mice per treatment group and stored frozen at −80°C prior to performing untargeted lipidomic analyses. For lipidomic analyses, each frozen hippocampus was weighed in a 1.5-mL safe-lock Eppendorf tube. The tube was added with two metal beads and 2 µL of water per mg of raw tissue. The samples were homogenized at a shaking frequency of 30 Hz on a MM 400 mill mixer for 1 min twice. Methanol-chloroform (3:1, v/v) at 18 µL per mg of raw tissue was then added. The samples were homogenized again for 1 min twice, followed by sonication in an ice-water bath for 3 min before centrifugal clarification at 21 000 g and 5°C for 10 min. The clear supernatants were quantitatively transferred to another set of Eppendorf tubes, where 240 µL of the clear supernatant was mixed with 120 µL of water-methanol (2:1 v/v) and 100 µL of chloroform. The mixture was vortex-mixed for 1 min at 3 000 rpm and then centrifuged to split the whole phase into an upper aqueous phase and a lower organic phase. The organic phase of each sample was carefully collected and dried under a nitrogen gas flow. The dried residue was dissolved in 120 µL of HPLC-grade ethanol. Aliquots of 6 µL from each solution were injected into a Waters BEH C4 LC column (2.1 I.D. * 50 mm, 1.7 μm for UPLC-high resolution mass spectrometry (HRMS) on a Thermo Ultimate 3000 UHPLC system coupled to a Thermo LTQ-Orbitrap Velos Pro mass spectrometer through an atmospheric pressure electrospray ionization (ESI) interface. The mobile phase was (A) 0.01% formic acid in water and (B) 0.01% formic acid in acetonitrile-isopropanol (1:1 v/v). The LC elution gradient was 5–50% B in 6 min; 50-100% B in 14 min and 100% B for 4 min, before the column was equilibrated at the initial solvent composition for 4 min between injections. The column flow rate was 400 μL/min and the column temperature was maintained at 40°C. Two UPLC-HRMS runs per sample were conducted in two rounds of LC injections with positive-ion and negative-ion detection, respectively. For lipid detection and relative quantitation, the MS instrument was operated with full-mass-Fourier transform MS detection, and at a mass resolution of 60 000 full width at half maximum (FWHM) at a mass-to-charge (*m/z*) ratio of 400. The mass range of HRMS detection was *m/z* ratios of 80 to 1800. A solution pooled from 20 μL aliquots of 12 randomly selected sample solutions was used as the quality control (QC) sample. This QC solution was injected at the beginning, in the middle and at the end of the UPLC-HRMS batch runs for each round of LC injections. During data acquisition, all the sample solutions were injected in a random order and two UPLC-HRMS datasets were acquired. Along with the UPLC-HRMS data acquisitions, LC-MS/MS of the QC sample solution was carried out using collision-induced dissociation (CID), with the 6 most abundant ions of each survey scan chosen for subsequent CID at normalized collision energies of 28–35%.

#### 2.7.2 Lipidomic data processing and analysis

The raw data files of UPLC-HRMS datasets with positive-ion and negative-ion detection were respectively processed using the XCMS module ^80–82^ in R with a custom-written script for peak detection, retention time shift correction, peak grouping and peak alignment in two rounds for each step. Mass deisotoping and removal of chemical background noise peaks were performed according to the seven golden rules described by ^83^. The output files of XCMS processing were in the format of *m/z*, retention time and peak area pair for each detected lipid, which was amenable for subsequent statistical analyses using MetaboAnalyst v5.0 ^84^. Prior to statistical analysis, the XCMS output data were quantile-normalized and log-transformed using the interquartile range (IQR) filtration and the detected lipid features detected in the QC samples, which showed relative standard deviations (RSDs) of >30% were removed. Group means were compared using two-sample t-test and fold-change (FC) analysis. Differentially-regulated lipids (DRLs) for each comparison were defined as having a false discovery rate (FDR)-adjusted p-value ≤0.05 and a fold change >1.5. DRLs were assigned by mass-matching against the Human Metabolome Database (HMDB) and LIPID MAPS databases within a maximum of mass errors of 5 ppm, in combination with spectral elucidation of the acquired lipid MS/MS spectra, with the aid of the MS/MS libraries of HMDB, METLIN, MASSBANK and LIPID MAPS and an in-house library of authentic compounds of different classes of >100 lipids. For lipid ontology enrichment analyses, LION software (v.2020.07.14) ^85^ was utilized. DRLs identified in the positive ion and negative ion detection modes were combined, and comparisons were made in ‘ranking mode’, using 2-log [fold change] analyses with a two-tailed alternative hypothesis in the K-S settings. Lipids that could not be matched to a LION identifier were retained in datasets during analysis.

### 2.8 Statistics

Data are reported as means ± standard error of the mean. Statistical analyses were conducted using Prism 9 (v.9.2, GraphPad Software, San Diego, CA, USA). Normality was verified using a Shapiro-Wilk test and assessed by QQ plot. All metabolic and immunological parameters were analyzed using a two-way analysis of variance (ANOVA) to compare diet (CD *versus* KD) and stress phenotypes (CTRL, SUS, or RES) as between-subject factors. Significant ANOVA tests with a main effect of diet or stress or a diet × stress interaction were reported. The different stress phenotypes (CTRL, SUS, or RES) were examined as different levels of the stress phenotype factor. Significant ANOVA tests with a main effect of either diet or stress or a diet × stress interaction were followed by Tukey *post-hoc* tests to identify significant differences between the relevant groups (CD CTRL *versus* KD CTRL, CD CTRL *versus* CD SUS, CD CTRL *versus* CD RES, CD SUS *versus* CD RES, KD CTRL *versus* KD SUS, KD CTRL *versus* KD RES, KD SUS *versus* KD RES, CD SUS *versus* KD SUS, CD RES *versus* KD RES). The differences were considered statistically significant with a *p* value < 0.05. Asterisks (*) were used to represent diet x stress interactions, hashtags (#) were used to represent results related to a main effect of stress, ampersands (&) were used to represent results related to a main effect of diet. Sample size (n) refers to individual animals for behavioral, metabolic, and molecular analyses. For morphology and ultrastructural analysis, n refers to individual microglia considering microglia as a biological unit, while N refers to the population size (number of animals). Analysis of microglia as a biological unit, instead of the animal, takes into account the high heterogeneity of the microglia population, allowing for the assessment of individual cellular contributions to the population response ^55,86,87^. No statistical outliers were removed.

## 3. Results

### 3.1 KD increases the proportion of stress-resistant mice following social stress

To study the mechanisms underlying the protective effects of a KD, we compared the outcomes of a KD *versus* CD in adult male mice exposed to RSD *versus* non-stressed CTRLs. Two-month-old C57BL/6J mice were introduced to KD or kept on CD starting 4 weeks prior to starting the paradigm until the end of the experiment (Fig. 1A). Ketosis was confirmed by measuring the levels of BHB from blood samples collected at Day 0 and Day 12 (Fig. 1B).

After 4 weeks, mice on both diets were exposed to 10 days of RSD (Days 1 to 10) while non-exposed mice served as CTRLs. The animals remained on their respective diet throughout the RSD protocol. To assess the susceptibility or resistance to stress, a SI test measuring social avoidance to a new CD1 mouse which is associated with stress susceptibility ^65^ was performed on Day 11. As described in details in the Methods, the social interaction arena was divided into CZ, an interaction and peri-interaction zones (Fig. 1A). A SI ratio was determined by dividing the time spent in the interaction zone and peri-interaction zone in the presence *versus* absence of a novel CD1 mouse. Mice with a SI ratio < 1 were classified as SUS and those with a ratio ≥ 1 as RES. The results were compared to paired-housed, non-stressed CTRL mice which received either CD or KD without exposure to social stress paradigm.

Blood levels of CORT were elevated in stressed animals thus confirming the effectiveness of RSD. Interestingly, when analyzing the SUS and RES separately, we also found an effect of stress (F(2, 40) = 7.933, *p*= 0.0013), showing increased levels of CORT in the SUS group compared to the CTRL and RES groups (Fig. 1C). SUS animals on both diets further displayed increased social avoidance compared to respective CTRLs as shown by their SI ratio (F(5, 65) = 18.75, *p*< 0.0001; CD CTRL 1.252 ± 0.2192 *versus* CD SUS 0.7184 ± 0.2187 *versus* CD RES 1.150 ± 0.1198; KD CTRL 1.169 ± 0.1805 *versus* KD SUS 0.6895 ± 0.2809 *versus* KD RES 1.096 ± 0.0848) (Fig. 1D). Quantification of the CZ time before *versus* after the introduction of a CD1 aggressor showed that SUS animals on both diets increased their presence in the CZ (Fig. 1E). SI ratio also revealed that animals following KD *versus* CD were more likely to be classified as RES (CD: 63.63% SUS: 36.36% RES; KD: 42.85% SUS: 57.14% RES) (Fig. 1F). We then performed a chi-square test, which showed a tendency for an increase in the proportion of mice classified as RES under the KD [chi-square, df: 2.131,1; z: 1.460; *p* = 0.0722] (Fig. 1F). The shift in the SUS:RES ratio observed in KD mice indicates a potential increase in stress resistance, compared to the expected proportion of 30% to 40% of RES using the same paradigm ^65,66^. Overall, these results support the idea that KD as a dietary intervention could confer stress protection and result in behavioral improvements. Our findings thus reveal that KD potentially increases resistance to RSD, by increasing the number of mice classified as RES.

The KD diet exerts anti-inflammatory properties, which could mediate its stress-resistance capacities ^5,8,11–13^. To better understand the mechanisms underlying the tendency for KD to have a protective effect, we performed multiplex ELISA and measured blood levels of 31 pro- and anti-inflammatory cytokines in mice exposed to RSD. To look at the effects of stress over time, our analyses were conducted on Day 6 and Day 12 of the RSD. On Day 6, there was a significant increase in 4 pro-inflammatory cytokines in the SUS animals *versus* CTRLs related to stress:, namely G-CSF, IL-6, IL-13 and IP-10 (G-CSF (F(1,14) = 4.253, #*p* = 0.0374), IL-6 (F(2,16) = 3.719, #*p* = 0.0472), IL-13 (F(2,16 = 8.230) ##*p* = 0.0035), IP-10 (F(2,16) = 4.181, #*p* = 0.0346) (Fig. 2A-E). On Day 12, only G-CSF presented a significant effect of stress, being significantly higher in the SUS *versus* CTRL and RES groups (F(2,16) = 7.723, ##*p* = 0.0045) (Fig. 2A-E’). A main effect of diet was observed in IFN-L on day 6 (F(1, 13 = 5.165) &*p* = 0.0407) (Fig. 2B) and on G-CSF on day 12 (F(1, 16) = 9.754 &&*p* = 0.0066) (Fig. 2A’).

**Figure 2:**
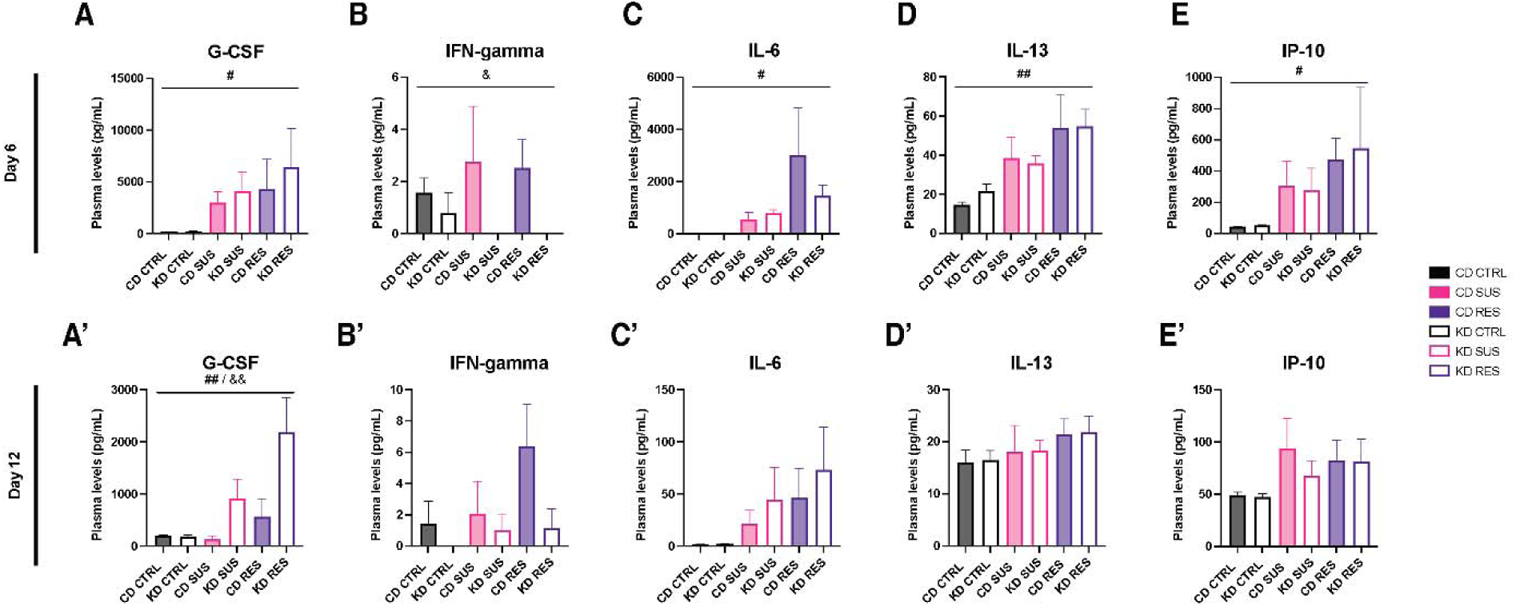
Ketogenic diet and repeated social defeat stress modify blood levels of inflammatory cytokines. On Day 6, granulocyte colony stimulating factor (G-CSF) **(A)**, interleukin 6 (IL-6) **(C)**, interleukin 13 (IL-13) **(D)** and C-X-C motif chemokine ligand 10 (IP-10) **(E)** presented an effect of stress. Interferon gamma (IFN-L) showed an effect of diet **(B)**. By Day 12, only G-CSF maintained changes related to stress **(A’)**. On Day 12, G-CSF showed an effect related to diet. n = 3–4 mice/group. Data are expressed as mean ± standard error of the mean. Statistical significance was assessed by 2-way ANOVA where #p < 0.05; ##p < 0.01 represent a main effect of stress; &p < 0.05; &&p < 0.01 represent a main effect of diet. CD: control diet; KD: ketogenic diet; CTRL: control; SUS: susceptible; RES: resistant; G-CSF: granulocyte colony stimulating factor; IFN-g: interferon-gamma; IL: interleukin; IP-10: C-X-C motif chemokine ligand 10.

Overall, our results indicate that 4 weeks of KD *versus* CD elevated the circulating levels of ketone bodies in mice. Ten days of RSD also caused behavioral changes in mice, resulting in distinct SUS and RES phenotypes. Ten days of RSD induced inflammation in the stressed mice, which differed in their circulating levels of inflammatory cytokines. Furthermore, the population following a KD showed a tendency toward an increased proportion of RES animals upon RSD

### 3.2 KD results in different microglial morphological adaptations to social stress

Microglia have emerged as key mediators of chronic stress outcomes by coordinating the peripheral and central immune responses to environmental challenges ^88–90^. To study whether changes in microglia might underlie the effects of KD at steady-state and upon psychosocial stress, we first analyzed possible changes in their density and distribution among the ventral hippocampus CA1 *stratum radiatum,* comparing the two diets, CD and KD, and three stress phenotypes, CTRL, SUS, and RES, on Day 12 of the paradigm. A double immunostaining for the microglia/macrophage marker Iba1 and largely microglia-specific TMEM119 was performed ^71^. Microglia were identified as Iba1-positive (+)/TMEM119+ cells (Fig. 3A). Microglial density (cells/area), nearest neighbor distance (NND; average distance of each microglia to its nearest neighbor) and spacing index (square average NND multiplied by the density) were assessed. This analysis revealed that the different parameters analyzed remained unchanged across all groups (Fig. 3B-D) (Table 1).

**Figure 3:**
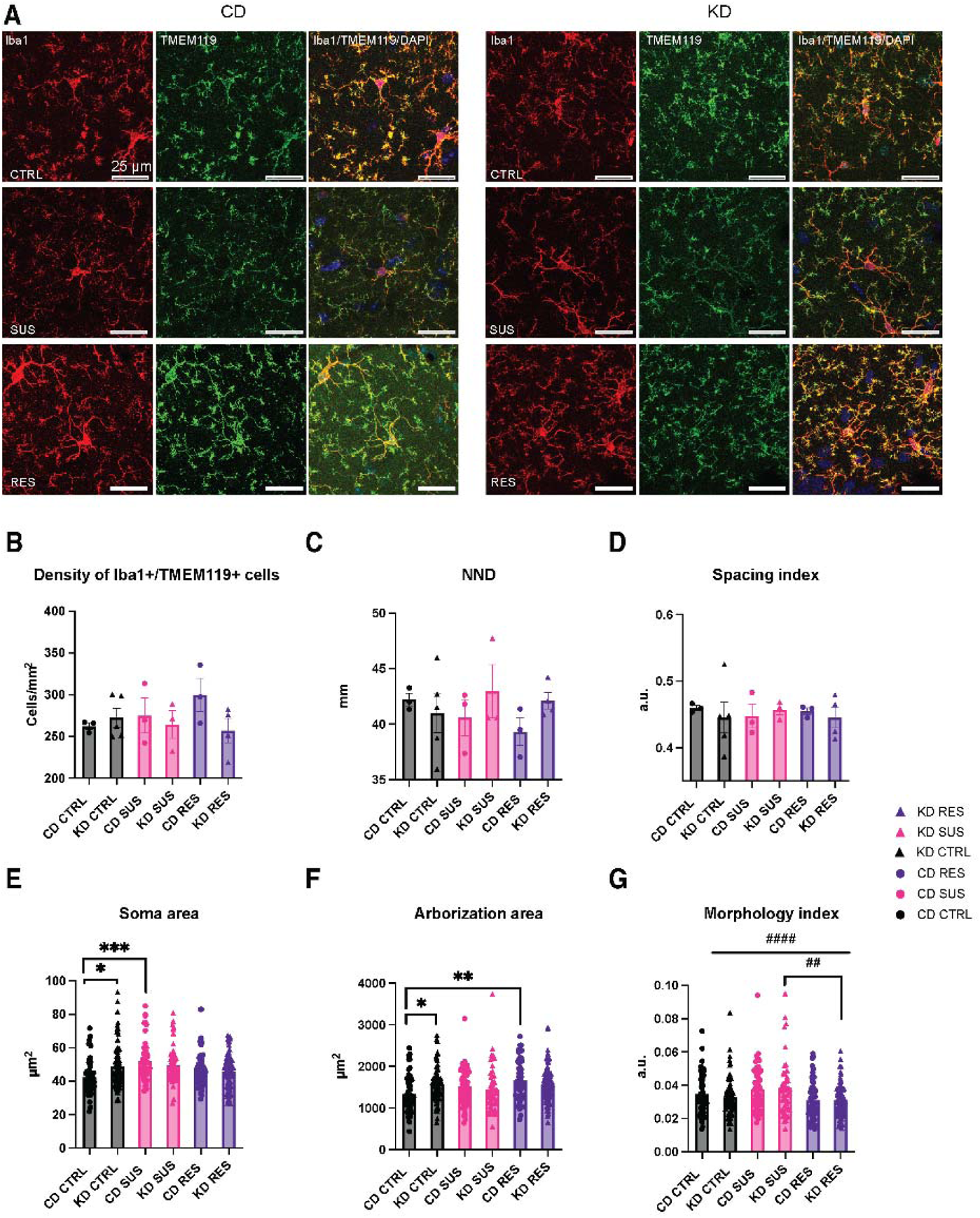
Ketogenic diet results in different microglial morphological adaptations to social stress. Representative confocal images at 40x magnification showing Iba1 (red) and TMEM119 (green) stained microglia in the ventral hippocampus CA1 *stratum radiatum* of the 6 experimental groups **(A)**. Scale bar is equivalent to 25 µm. Microglial density and distribution remained unchanged by stress or diet as observed by density **(B)**, nearest neighbor distance (NND) **(C)** and spacing index **(D)** of Iba1+/TMEM119+ cells. Microglia of KD fed mice showed an increase in soma **(E)** and arborization **(F)** area in non-stressed controls (CTRL). Microglia of stressed animals showed morphological adaptations to stress: an increase of soma area **(E)** of control diet (CD) susceptible (SUS) microglia compared to CD CTRL, and an increase in arborization **(F)** area of CD resistant (RES) microglia compared to CD CTRL. Stress modified the microglial morphology index **(G)**. Microglia of SUS animals had a bigger soma to arborization ratio compared to CTRL and RES animals of KD group. n = 51–88 microglia/group; N = 3–5 mice/group. Data are expressed as mean ± standard error of the mean. Statistical significance was assessed by 2-way ANOVA followed by Tukey *post-hoc* analysis, where *p < 0.05, **p < 0.01, ***p < 0.001. ##p < 0.01, ####p < 0.0001 represent a main effect of stress. CD: control diet; KD: ketogenic diet; CTRL: control; SUS: susceptible; RES: resistant; NND: nearest neighbor distance; a.u.: arbitrary units.

**Table 1.**
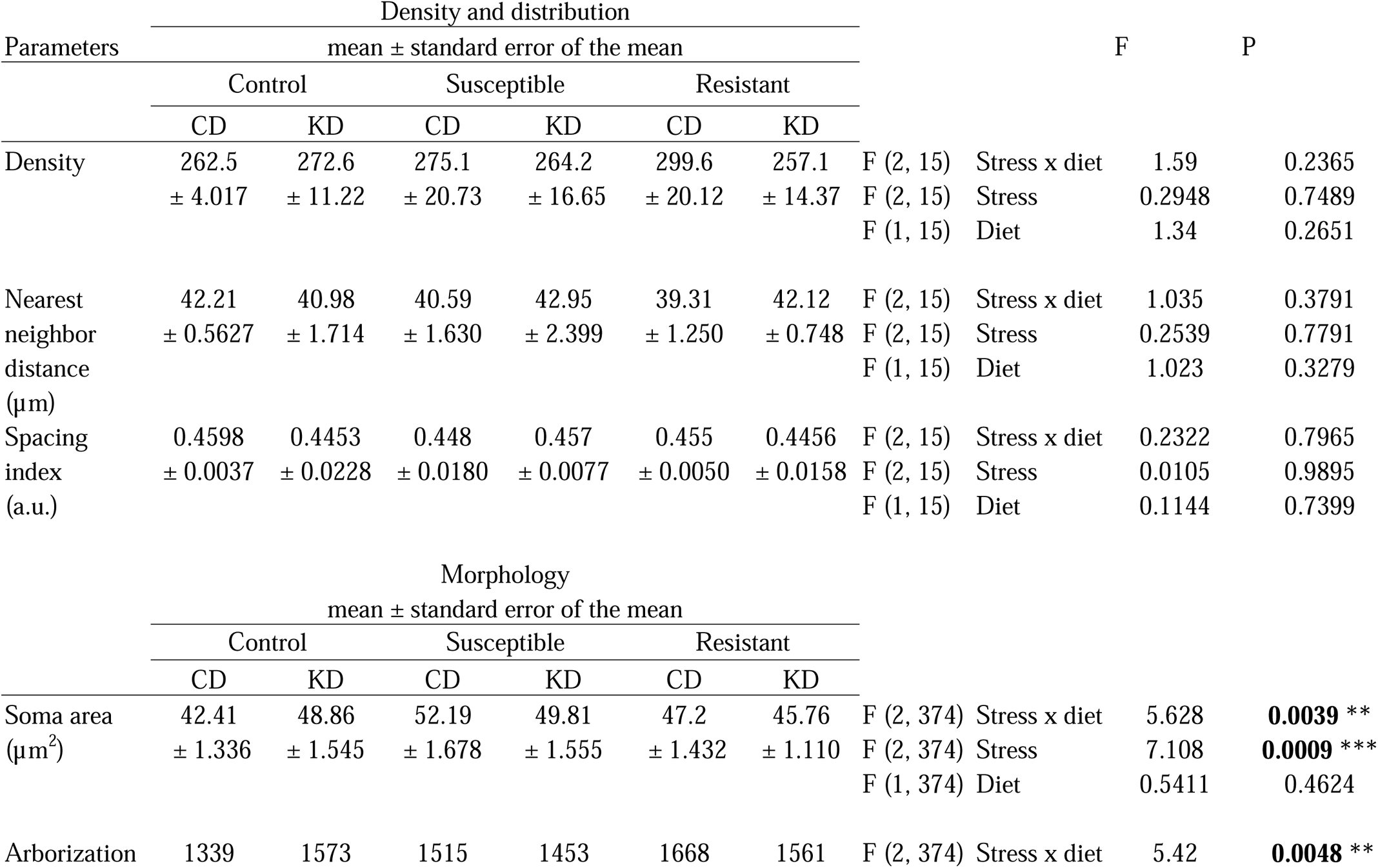
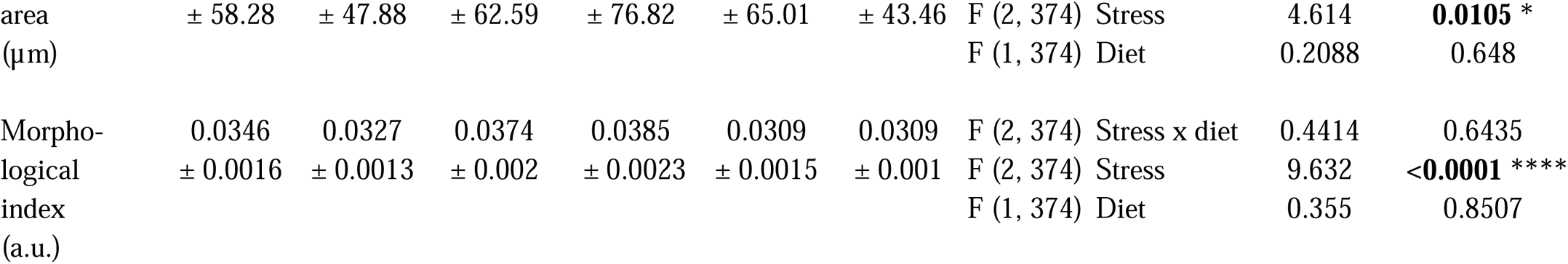
Microglial number, distribution and morphological properties in CD *versus* KD fed mice exposed to RSD. a.u.: arbitrary units; CD: control diet; KD: ketogenic diet; RSD: repeated social defeat.

| Parameters | Density and distribution |  |  |  |  |  |  |  | F | P |
| --- | --- | --- | --- | --- | --- | --- | --- | --- | --- | --- |
|  | mean ± standard error of the mean |  |  |  |  |  |  |  |  |  |
|  | Control |  | Susceptible |  | Resistant |  |  |  |  |  |
|  | CD | KD | CD | KD | CD | KD |  |  |  |  |
| Density | 262.5 | 272.6 | 275.1 | 264.2 | 299.6 | 257.1 | F (2, 15) | Stress x diet | 1.59 | 0.2365 |
|  | ± 4.017 | ± 11.22 | ± 20.73 | ± 16.65 | ± 20.12 | ± 14.37 | F (2, 15) | Stress | 0.2948 | 0.7489 |
|  |  |  |  |  |  |  | F (1, 15) | Diet | 1.34 | 0.2651 |
| Nearest neighbor distance (µm) | 42.21 | 40.98 | 40.59 | 42.95 | 39.31 | 42.12 | F (2, 15) | Stress x diet | 1.035 | 0.3791 |
|  | ± 0.5627 | ± 1.714 | ± 1.630 | ± 2.399 | ± 1.250 | ± 0.748 | F (2, 15) | Stress | 0.2539 | 0.7791 |
|  |  |  |  |  |  |  | F (1, 15) | Diet | 1.023 | 0.3279 |
| Spacing index (a.u.) | 0.4598 | 0.4453 | 0.448 | 0.457 | 0.455 | 0.4456 | F (2, 15) | Stress x diet | 0.2322 | 0.7965 |
|  | ± 0.0037 | ± 0.0228 | ± 0.0180 | ± 0.0077 | ± 0.0050 | ± 0.0158 | F (2, 15) | Stress | 0.0105 | 0.9895 |
|  |  |  |  |  |  |  | F (1, 15) | Diet | 0.1144 | 0.7399 |
|  | Morphology |  |  |  |  |  |  |  |  |  |
|  | mean ± standard error of the mean |  |  |  |  |  |  |  |  |  |
|  | Control |  | Susceptible |  | Resistant |  |  |  |  |  |
|  | CD | KD | CD | KD | CD | KD |  |  |  |  |
| Soma area (µm<sup>2</sup>) | 42.41 | 48.86 | 52.19 | 49.81 | 47.2 | 45.76 | F (2, 374) | Stress x diet | 5.628 | **0.0039 \*\*** |
|  | ± 1.336 | ± 1.545 | ± 1.678 | ± 1.555 | ± 1.432 | ± 1.110 | F (2, 374) | Stress | 7.108 | **0.0009 \*\*\*** |
|  |  |  |  |  |  |  | F (1, 374) | Diet | 0.5411 | 0.4624 |
| Arborization | 1339 | 1573 | 1515 | 1453 | 1668 | 1561 | F (2, 374) | Stress x diet | 5.42 | **0.0048 \*\*** |

|  |  |  |  |  |  |  |  |  |  |
| --- | --- | --- | --- | --- | --- | --- | --- | --- | --- |
| area<br>($\mu\text{m}$) | $\pm 58.28$ | $\pm 47.88$ | $\pm 62.59$ | $\pm 76.82$ | $\pm 65.01$ | $\pm 43.46$ | F (2, 374) Stress<br>F (1, 374) Diet | 4.614<br>0.2088 | **0.0105 \***<br>0.648 |
| Morpho-<br>logical<br>index<br>(a.u.) | 0.0346<br>$\pm 0.0016$ | 0.0327<br>$\pm 0.0013$ | 0.0374<br>$\pm 0.002$ | 0.0385<br>$\pm 0.0023$ | 0.0309<br>$\pm 0.0015$ | 0.0309<br>$\pm 0.001$ | F (2, 374) Stress x diet<br>F (2, 374) Stress<br>F (1, 374) Diet | 0.4414<br>9.632<br>0.355 | 0.6435<br>**<0.0001 \*\*\*\***<br>0.8507 |

We then assessed in the same animals and hippocampal region, also on Day 12, possible changes in microglial morphology under the two diets (Table 1). We found in the KD-fed mouse population that microglia had morphological differences at steady-state, among the non-stressed CTRL group (Table 1). Two-way ANOVA showed changes in soma (F(2,374) = 5.628 \*\**p* = 0.0039) and *post-hoc* analysis indicated that microglia had a bigger soma in KD CTRL compared to CD CTRL mice (CD CTRL 42.41 µm^2^ ± 1.336 *versus* KD CTRL 48.86 µm^2^ ± 1.545 \**p* =0.0169). Two-way ANOVA revealed a difference in arborization area (F(2,374) = 5.42 \*\**p* = 0.0048) and *post-hoc* analysis indicated that microglia of KD CTRL *versus* CD CTRL animals had a larger arborization area (CD CTRL 1339 µm^2^ ± 58.28 *versus* KD CTRL 1573 µm^2^ ± 47.88 \**p* =0.0391). These differences in microglial soma and arborization area observed in unstressed CTRL animals following a KD indicate that this diet modifies microglial properties at the basal level, under steady-state conditions.

We further examined possible changes in microglial morphology induced by stress under the two diets (Table 1).Two-way ANOVA revealed an effect of stress (F(2, 374) = 7.108 \**p* = 0.0009) and *post-hoc* analysis indicated that microglia in the CD SUS group had increased soma area compared to the CD CTRL group (CD CTRL 42.41 µm^2^ ± 1.336 *versus* CD SUS 52.19 µm^2^ ± 1.678 \*\*\**p* =0.0002) (Fig. 3E). Microglia in CD RES mice had an increased arborization area compared to those in CD CTRL animals (CD CTRL 1339 µm^2^ ± 58.28 *versus* CD RES 1668 µm^2^ ± 65.01 \*\**p* = 0.0017) (Fig. 3E). These morphological changes were not observed upon stress in KD animals, for soma area (KD CTRL 48.86 µm^2^ ± 1.545 *versus* KD SUS 49.81 µm^2^ ± 1.555) and arborization area (KD CTRL 1573 µm^2^ ± 47.88 *versus* KD RES 1561 µm^2^ ± 43.46) (Fig. 3E-F). A main effect of stress (F(2,374) = 9.632, ####*p* < 0.0001) was, however, observed for the morphology index. *Post-hoc* analysis revealed that microglia in KD RES *versus* KD SUS mice had a smaller morphology index (KD SUS 0.03852 a.u. ± 0.0023 *versus* KD RES 0.3094 a.u. ± 0.0010 ##*p* = 0.0066) (Fig. 3G), meaning that microglia have a smaller soma in relationship to their arborization, a ratio that describes more ramified microglia. While changes in microglial morphology can provide relevant insights into their physiological functions ^91,92^, these results indicate that microglia display diverse morphological adaptations to diet and stress.

### 3.3 KD changes microglia-synapse interactions differently in stress susceptibility and resistance

To characterize how KD and RSD affect microglial interactions within the hippocampus, a detailed ultrastructural analysis was performed with SEM in the CA1 *stratum radiatum* of the same animals, also on Day 12. Microglial contacts with neuronal and astrocytic cell bodies as well as blood vessels (basement membrane) were first quantified. There were no changes in these microglial interactions between CTRL, SUS, and RES mice under both diets (Table 2).

**Table 2.**
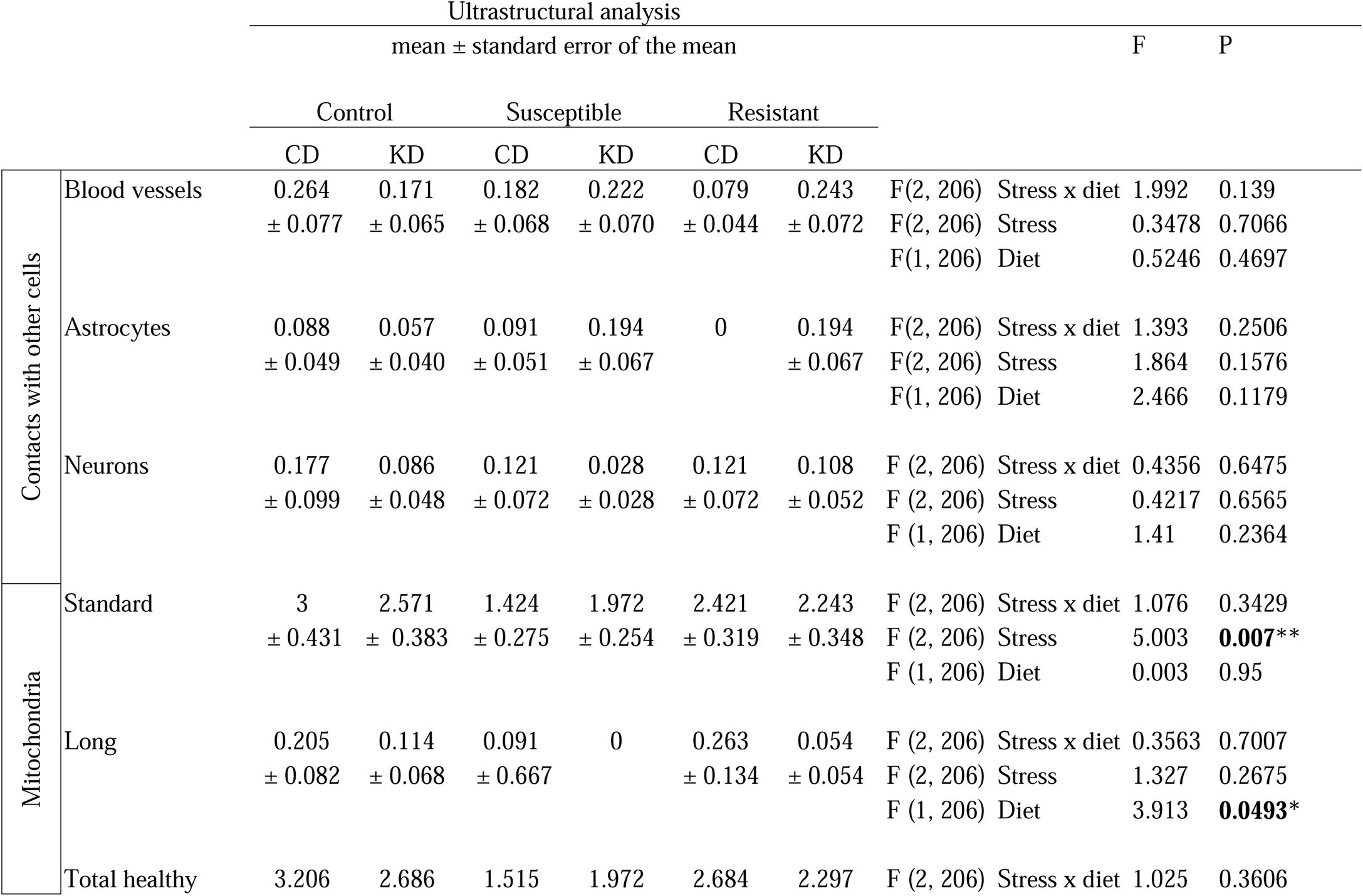
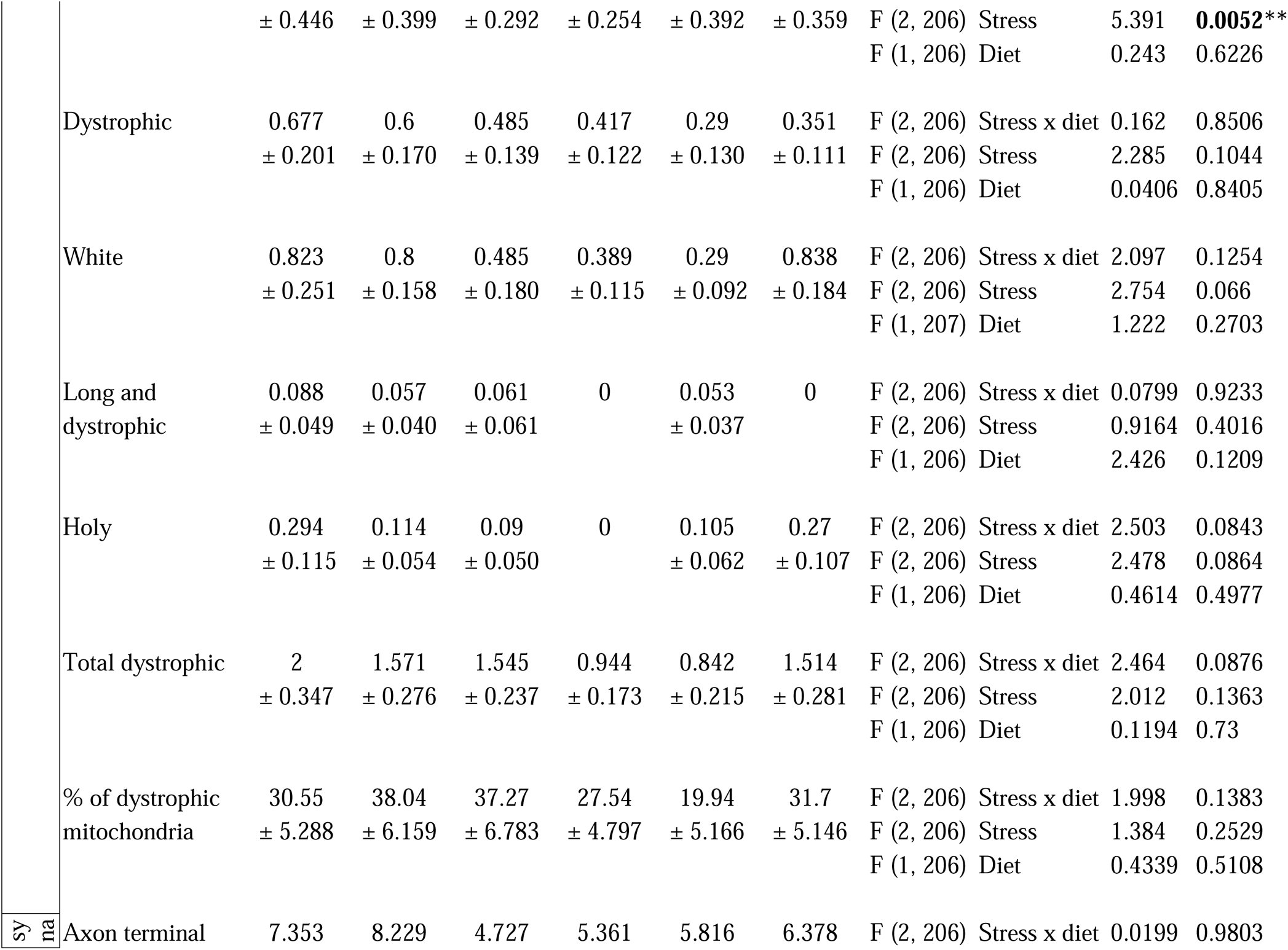
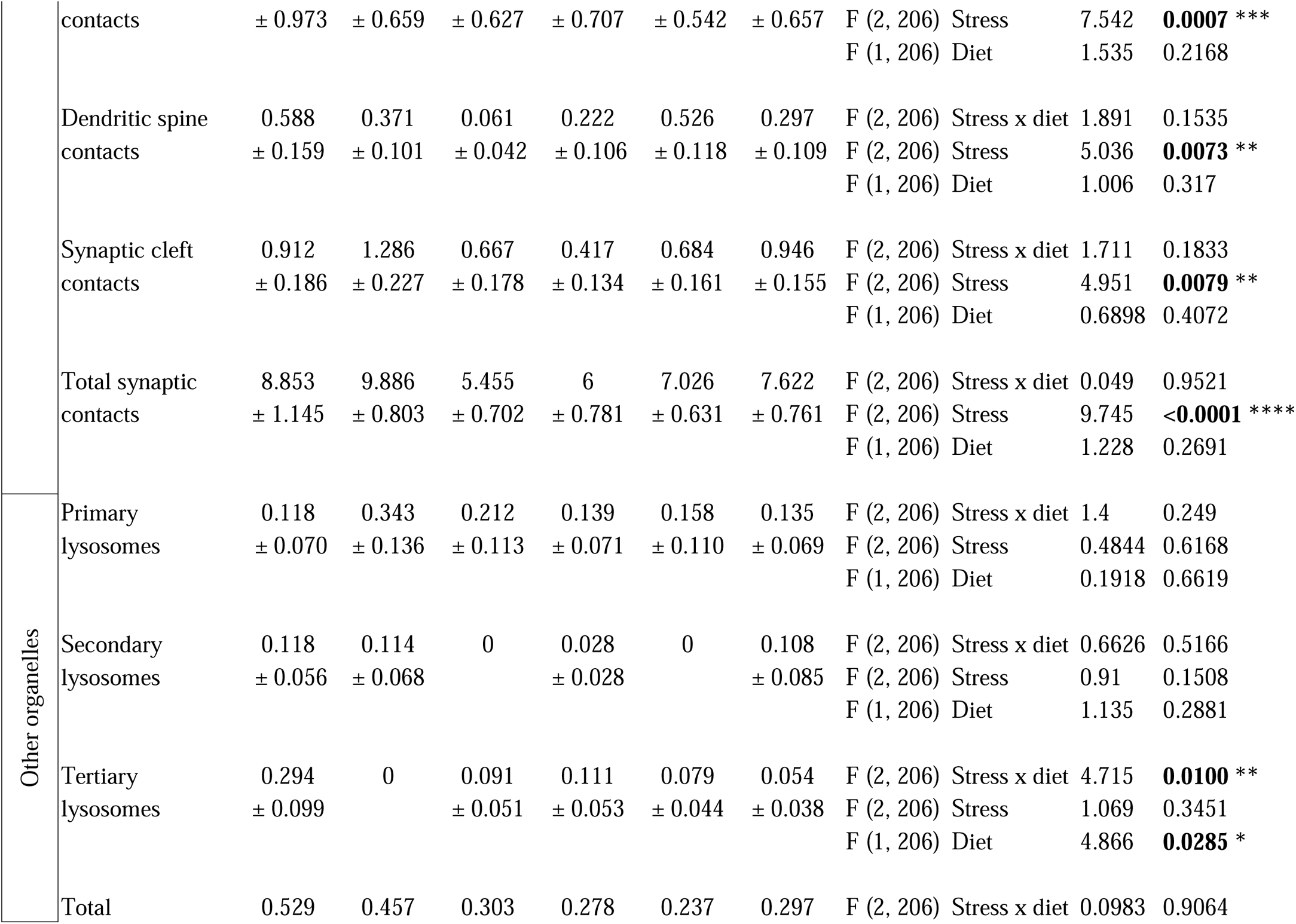
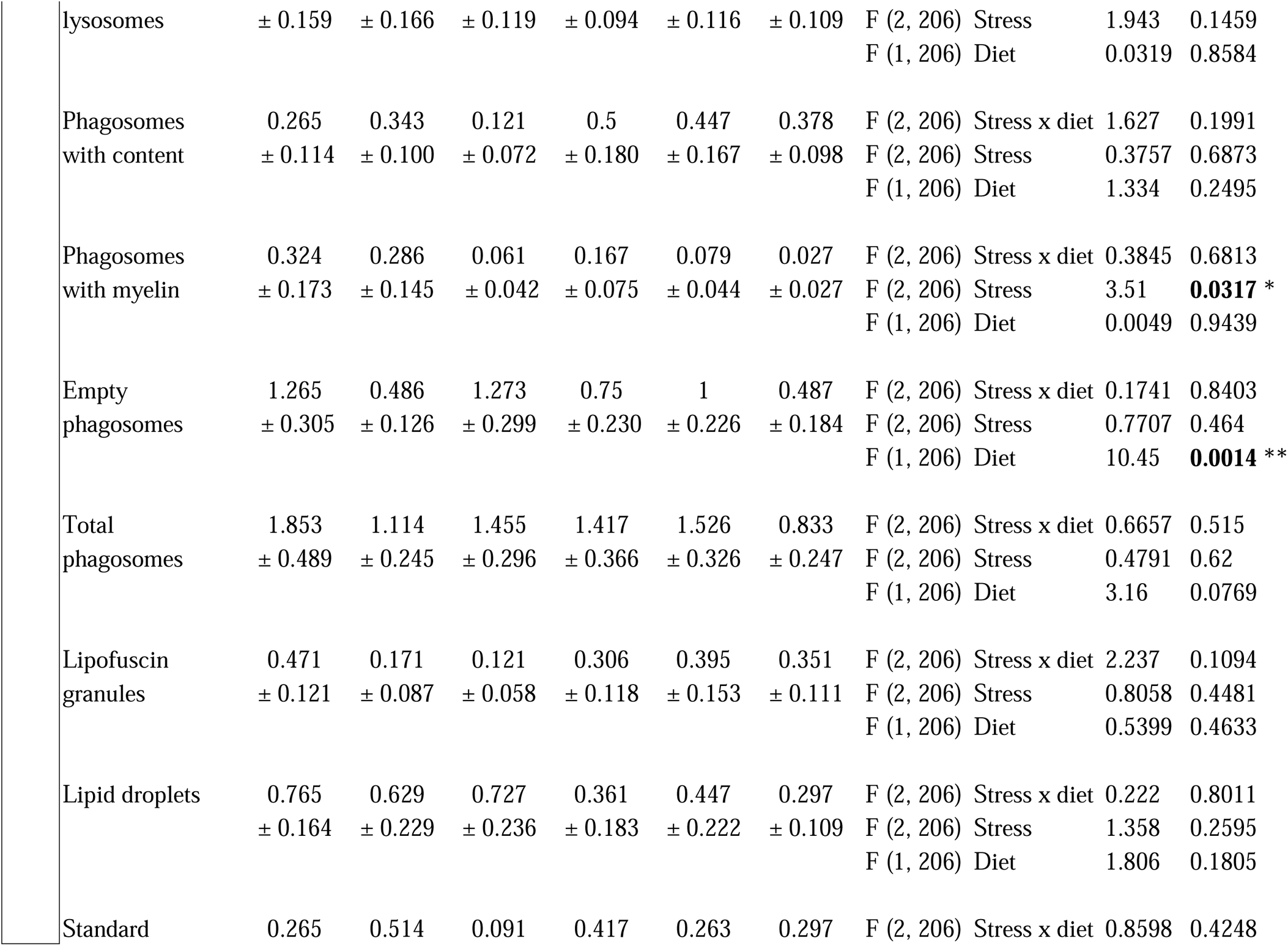
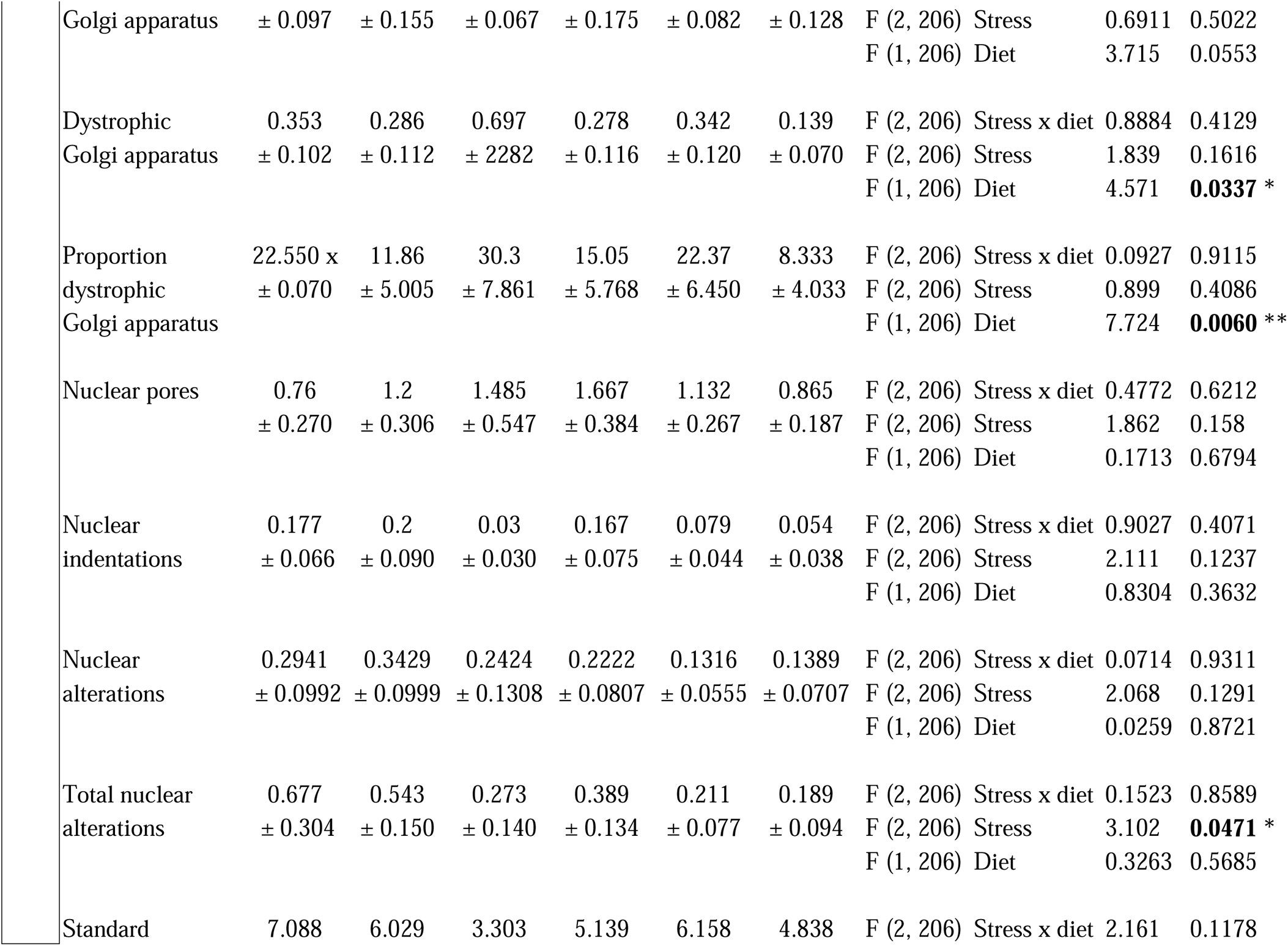
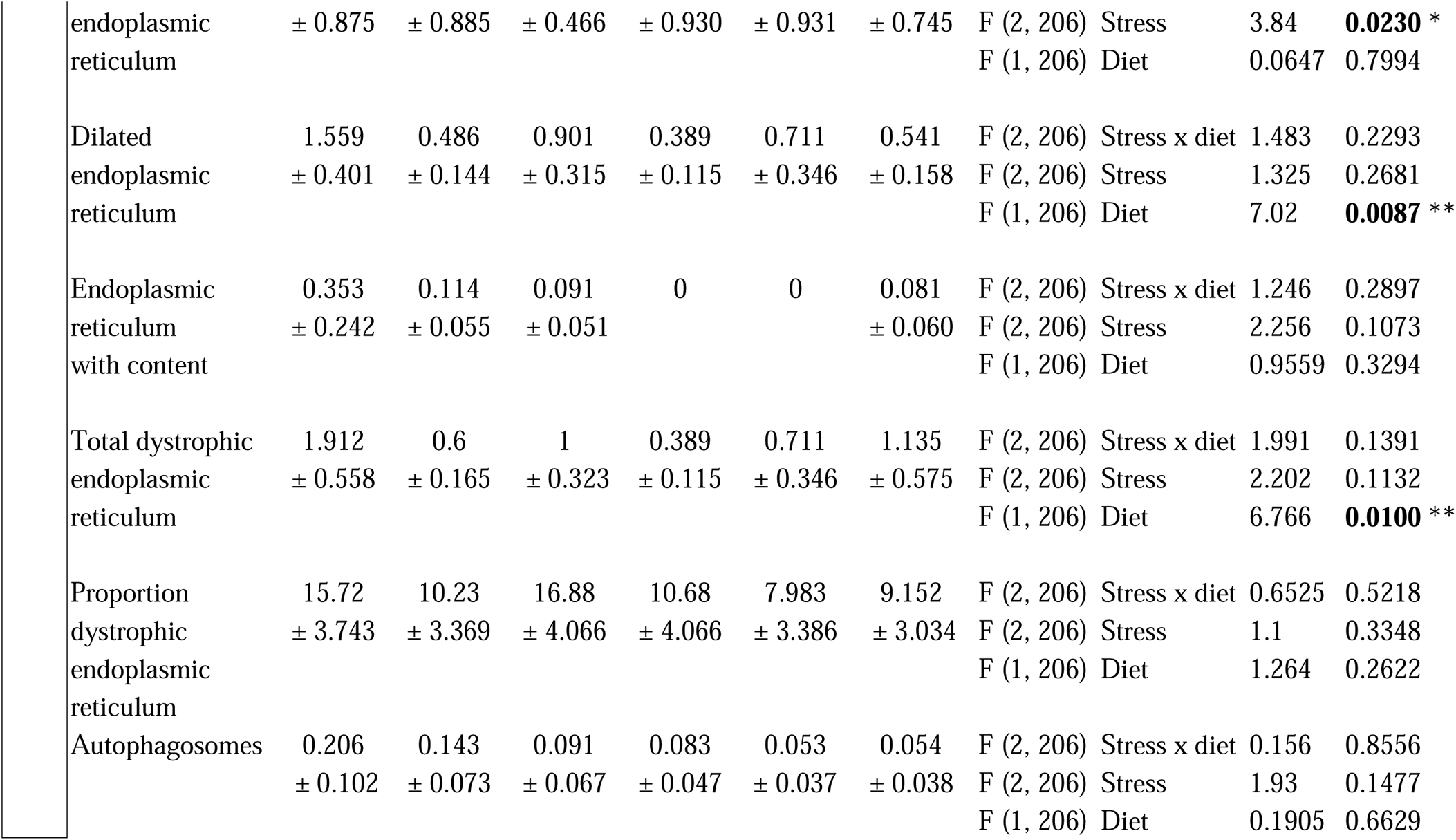
Microglial ultrastructural features in CD *versus* KD fed mice exposed to RSD. CD: control diet; KD: ketogenic diet; RSD: repeated social defeat.

As microglia-synapse interactions are relevant to the regulation of neuronal activity and plasticity processes, notably upon stress and in MDD ^93^, microglial direct contacts with excitatory synapses were next quantified. They were classified as contacts with presynaptic axon terminals (identified by their synaptic vesicles), dendritic spines (recognized by their post-synaptic density) and synaptic clefts (contact with both pre- and post-synaptic elements) (Fig. 4A-D). Diet had no effect on microglial contacts with synaptic elements between the groups (Fig. 4E-H) (Table 2). However, we observed a main effect of stress on the number of direct microglial contacts with axon terminals (F(2, 206) = 7.542, ###*p* = 0.0007), dendritic spines (F(2, 206) = 5.036, ##*p* = 0.0073) and synaptic clefts (F(2, 206) = 4.951, ##*p* = 0.0079) (Fig. 4E-G). *Post-hoc* analysis further revealed that the CD SUS mice had a reduced number of microglial contacts with dendritic spines than CD CTRL and CD RES (CD CTRL 0.5882 ± 0.1586 *versus* CD SUS 0.0606 ± 0.0421 #*p* < 0.0174 *versus* CD RES 0.526 ± 0.118 #*p* = 0.0426) (Fig. 4F). KD SUS microglia also showed fewer contacts than KD CTRL with axon terminals (KD CTRL 8.229 ± 0.6586 *versus* KD SUS 5.361 ± 0.7073 #*p* = 0.0486) and synaptic clefts (KD CTRL 1.286 ± 0.2267 *versus* KD SUS 0.4167 ± 0.1344 ##*p* = 0.007) (Fig. 4E-G). To assess the overall impact of stress on microglial interventions at neuronal circuits, when synaptic element types were pooled, 2-way ANOVA analysis showed a main effect of stress (F(2, 206) = 9.745, ####*p* < 0.0001) on the total number of microglial contacts with synaptic elements (Fig. 4H). Following *post-hoc* analysis a decrease in the number of these microglial contacts with synaptic elements between KD SUS and KD CTRL groups (KD CTRL 9.886 ± 0.8031 *versus* KD SUS 6.0 ± 0.7807 #*p* = 0.0113) was observed (Fig. 4H).

**Figure 4:**
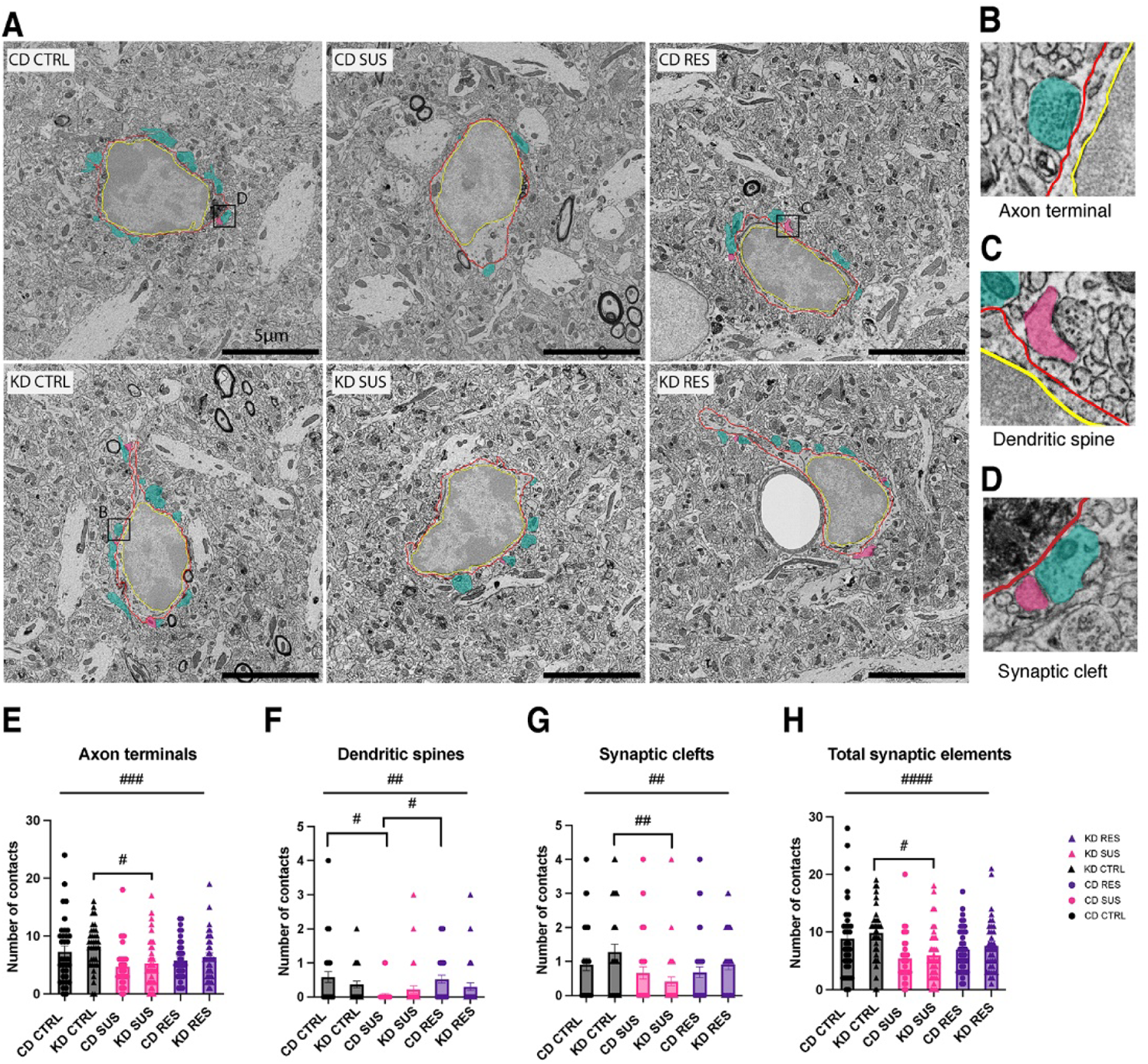
Microglia-synapse interactions change differently in stress susceptibility and resistance. Representative 5 nm resolution scanning electron microscopy images showing microglia captured in the ventral hippocampus CA1 *stratum radiatum* of the 6 experimental groups **(A)**. There is a reduction in the number of direct contacts between microglial cell bodies and pre-synaptic axon terminals **(B,F)**, post-synaptic dendritic spines **(C,G)** and simultaneous contact to two synapse-forming elements classified as synaptic clefts **(D,H)**. Stress affects plasticity as reflected by total number of contacts between microglial cell bodies and all three categories of synaptic elements **(I)**. n = 34–38 cells/group; N = 3 mice/group. Data are expressed as mean ± standard error of the mean. Statistical significance was assessed by 2-way ANOVA followed by Tukey *post-hoc* analysis, where #p < 0.05, ##p < 0.01, ###*p < 0.001,* ####p < 0.0001 represent a main effect of stress. CD: control diet; KD: ketogenic diet; CTRL: control; SUS: susceptible; RES: resistant; red outline: microglial plasma membrane; yellow outline: nuclear membrane; green pseudo-coloring: pre-synaptic axon terminals; magenta pseudo-coloring: post-synaptic dendritic spines.

Together, these results reveal that SUS mice display reduced microglial contacts with synaptic elements. Increasing evidence has revealed the importance of membrane-to-membrane contacts between microglia and their neighboring cells in shaping neuronal networks, for instance via neuronal synchronization, synaptic plasticity, and structural remodeling ^94,95^.

### 3.4 KD and stress differently modify microglial phagolysosomal inclusions and other organelles

To provide further insights into possible changes in microglial function with KD and RSD, we next examined their accumulation of cellular stress markers, indicative of altered or compromised activities ^73^. We performed an ultrastructural characterization of microglial organelles as well as classified and quantified their markers of cellular stress in samples from the same animals and region imaged by SEM on Day 12. We examined microglial mitochondria (healthy, long, altered), lysosomal inclusions (primary, secondary, tertiary lysosomes), autophagosomes, phagosomes (with content or empty), lipofuscin granules, lipid droplets, ER and Golgi apparatus (standard or with alterations), and nucleus (pores, indentations, alterations).

We detected an interaction of diet x stress for the number of tertiary lysosomes (F(2,206) = 4.715, \*\**p* < 0.01) (Fig. 5A-C). *Post-hoc* analysis further showed that microglia in CD CTRL *versus* KD CTRL mice had more tertiary lysosomes (CD CTRL 0.294 ± 0.099 *versus* KD CTRL 0 \*\**p* = 0.0032) (Fig. 5C). We observed a main effect of stress on a series of ultrastructural characteristics associated with phagocytic activity as well as cellular stress and aging (Table 2). In particular, we found a main effect of stress on the number of healthy ER (F(2, 206) = 3.840 #*p* = 0.0230) (Fig. 5G). *Post-hoc* analysis revealed that CD SUS animals compared to CD CTRL had a reduced number of healthy ER (CD CTRL 7.088 ± 0.8748 *versus* CD SUS 3.303 ± 0.4656 #*p* = 0.0247) (Fig. 5G). We further observed diet-related ultrastructural changes associated to cellular stress in ER and Golgi apparatus (Fig. 5D-F). Two-way ANOVA analysis revealed a main effect of the diet, notably for the number of microglial ER with dilation (F(1,206) = 7.020 &&*p* = 0.0087) (Fig. 5H). Two-way ANOVA analysis revealed a main effect of the diet, notably for the total number of microglial ER with dystrophy (F(1,206) = 7.020 &&*p* = 0.0087) (Fig. 5I). *Post-hoc* analysis indicated that microglia in KD CTRL *versus* CD CTRL mice had a reduced number of ER showing dystrophy (CD CTRL 1.912 ± 0.5575 *versus* KD CTRL 0.600 ± 0.1650 &*p* = 0.0431) (Fig. 5I). Other features presenting a significant effect of the diet included the number of dilated Golgi apparatus (F(1,206) = 6.766 &*p* = 0.0337) (Fig. 5J), the proportion of dilated Golgi (F(1,206) = 7.724 &&*p* = 0.006) (Fig. 5K), the number of elongated mitochondria (F(1,206) = 3.913 &*p* = 0.0493) (Fig. 5L), and the number of empty phagosomes (F(1,206) = 10.45 &&*p* = 0.0014) (Fig. 5M).

**Figure 5:**
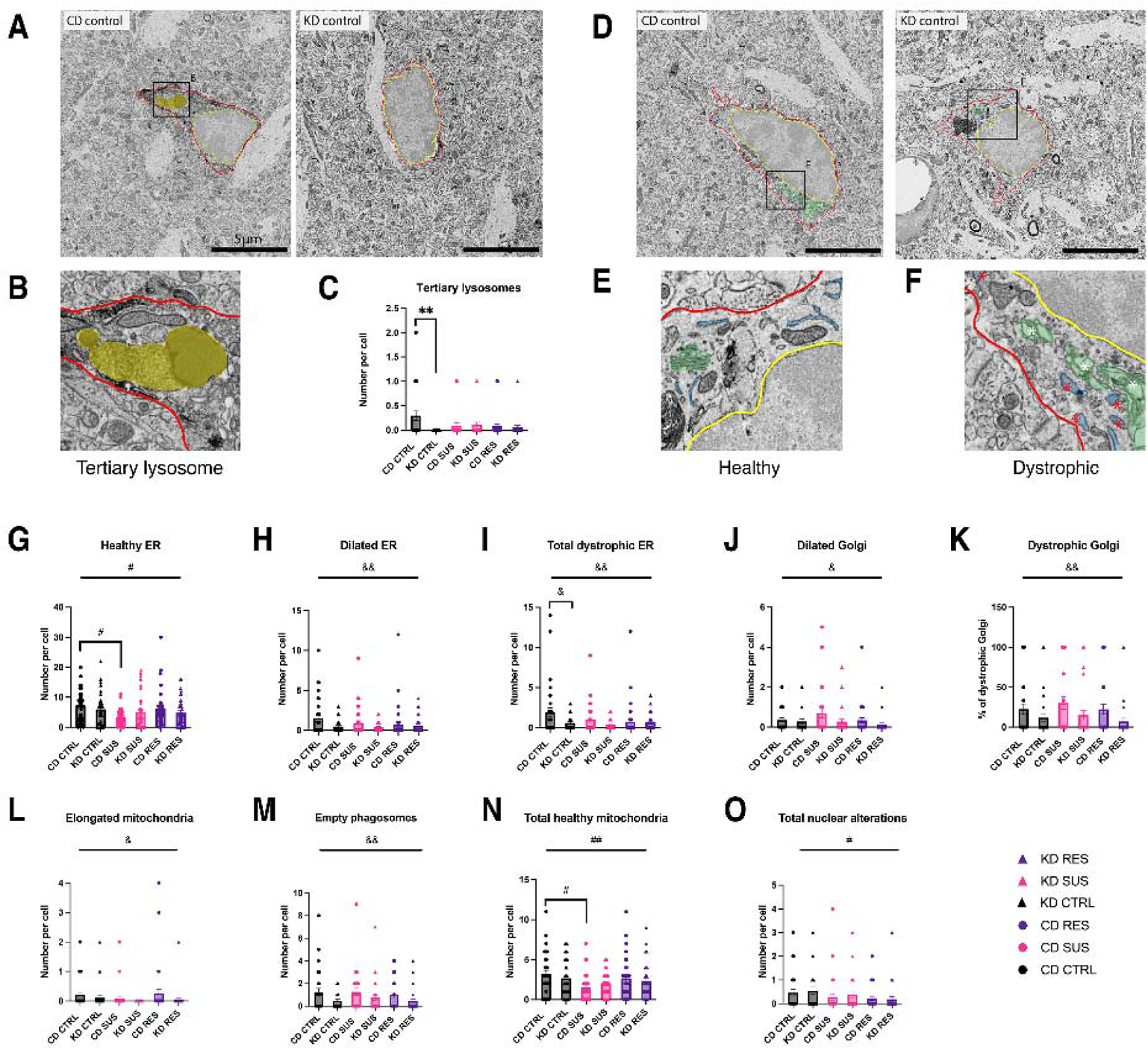
Ketogenic diet modifies microglial organelle number and ultrastructure. Representative 5 nm resolution scanning electron microscopy images showing microglia captured in the ventral hippocampus CA1 *stratum radiatum* of unstressed control animals **(A)**. Ultrastructural example of a tertiary lysosome **(B)**. There is a reduction in the number of tertiary lysosomes in microglial cell bodies from KD fed control animals compared to control diet (CD) fed controls **(C)**. Representative 5 nm resolution scanning electron microscopy images showing microglia captured in the ventral hippocampus CA1 *stratum radiatum* of non-stressed controls (CTRL). Ultrastructural examples of healthy and dystrophic endoplasmic reticulum (ER) and Golgi apparatus **(E, F)**. KD diet modifies in microglial cell bodies the number and ultrastructural characteristics of ER as seen by the total number of healthy ER **(G)**, ER showing signs of dilation **(H)** and total number of ER with signs of dystrophy (endoplasmic reticulum with inclusions and dilation) **(I)**. KD diet affects in microglial cell bodies the number and ultrastructural features of Golgi apparatus, as seen by changes in the number of dilated Golgi apparatus **(J)** and total number of Golgi apparatus with signs of dystrophy (Golgi apparatus with inclusions and dilation) **(K)**. Diet affects in microglial cell bodies the number of elongated mitochondria **(L)** and empty phagosomes **(M)**. Stress affects in microglial cell bodies the number of total healthy mitochondria (standard and long mitochondria with absence of dystrophy), **(N)** elongated mitochondria and total nuclear alterations **(O)**. n = 34–38 cells/group; N = 3 mice/group. Data are expressed as mean ± standard error of the mean. Statistical significance was assessed by 2-way ANOVA followed by Tukey *post-hoc* analysis, where **p < 0.01. #p < 0.05, ##p < 0.01 represent a main effect of stress. &p < 0.05, &&p < 0.01 represent a main effect of diet. CD: control diet; KD: ketogenic diet; CTRL: control; SUS: susceptible; RES: resistant; ER: endoplasmic reticulum. Red outline: microglial plasma membrane; yellow outline: nuclear membrane; yellow pseudo-coloring: tertiary lysosome; green pseudo-coloring: Golgi apparatus; blue pseudo-coloring: endoplasmic reticulum; white star dilated Golgi apparatus; red star: dilated ER.

We further observed a main effect of stress on mitochondria and the nuclear envelope. Total number of healthy mitochondria showed a main effect of stress (Fig. 5K) (F(2,206) = 5.391 ##*p* = 0.0052) (Fig. 5N). *Post-hoc* analysis revealed that CD SUS compared to CD CTRL had a reduced number of healthy mitochondria (CD CTRL 3.206 ± 0.4464 *versus* CD SUS 1.515 ± 0.2923 #*p* < 0.0205) (Fig. 5N). Additionally, we observed a main effect of stress on the number of nuclear alterations, which include indentations and alterations to the nuclear envelope (F(2,206) = 3.102 #*p* = 0.0471) (Fig. 5O).

These changes in ultrastructural features reveal that a KD diet and stress differently influence microglial properties, including phagolysosomal activity as well as cellular stress and aging.

### 3.5 KD alters the hippocampal lipidomic profile at steady-state and upon social stress

To deepen our understanding of the effects of KD on the hippocampal metabolism and function, we next performed lipidomic analyses of diet-related differences between mice with a shared stress phenotype (CTR, SUS or RES). Datasets from positive and negative ion data acquisition modes were analyzed separately for initial statistical investigations. Principal component analysis (PCA) demonstrated differential clustering of KD and CD samples (Fig. 6A). In CTRL mice, the major principal component (PC1) explained 22.5% and 26.2% of the variation among groups in the negative ion and positive ion datasets, respectively. Similarly, PC1 explained 26.4% and 28.3% of the variation in the SUS group, and 21.3% and 22.6% in the RES group, in negative ion and positive ion datasets, respectively. Two-sample t-tests and fold change analyses further revealed substantial numbers of differentially-regulated lipids (DRLs) between the KD and CD groups (Fig. 6B, Supplementary Figure 1A, Supplementary Table 1). The DRLs were subsequently annotated by matching to the Human Metabolome Database (HMDB), combining the positive and negative ion mode datasets, which led to successful annotation and identification of 32 DRLs between CTRL KD and CD mice, 21 DRLs between RES KD and CD mice, and 52 DRLs between SUS KD and CD mice. In response to the KD regime, hippocampal lipids belonging to several classes were highly upregulated (Fig. 6B, Supplementary Figure 1A, Supplementary Table 1). This included multiple phospholipids, such as phosphatidylglycerols, sphingomyelins, lysophosphatidylethanolamines (LPEs) and phosphatidylserines (PSs), with DRL profiles differing distinctly at the lipid species level among CTRL, SUS and RES mice in response to a KD. Several phospholipid classes were also downregulated in response to a KD in CTRL and SUS mice, whereas certain conjugated N-acyl taurines were uniquely downregulated in RES mice (Supplementary Figure 1A, Supplementary Table 1).

**Figure 6:**
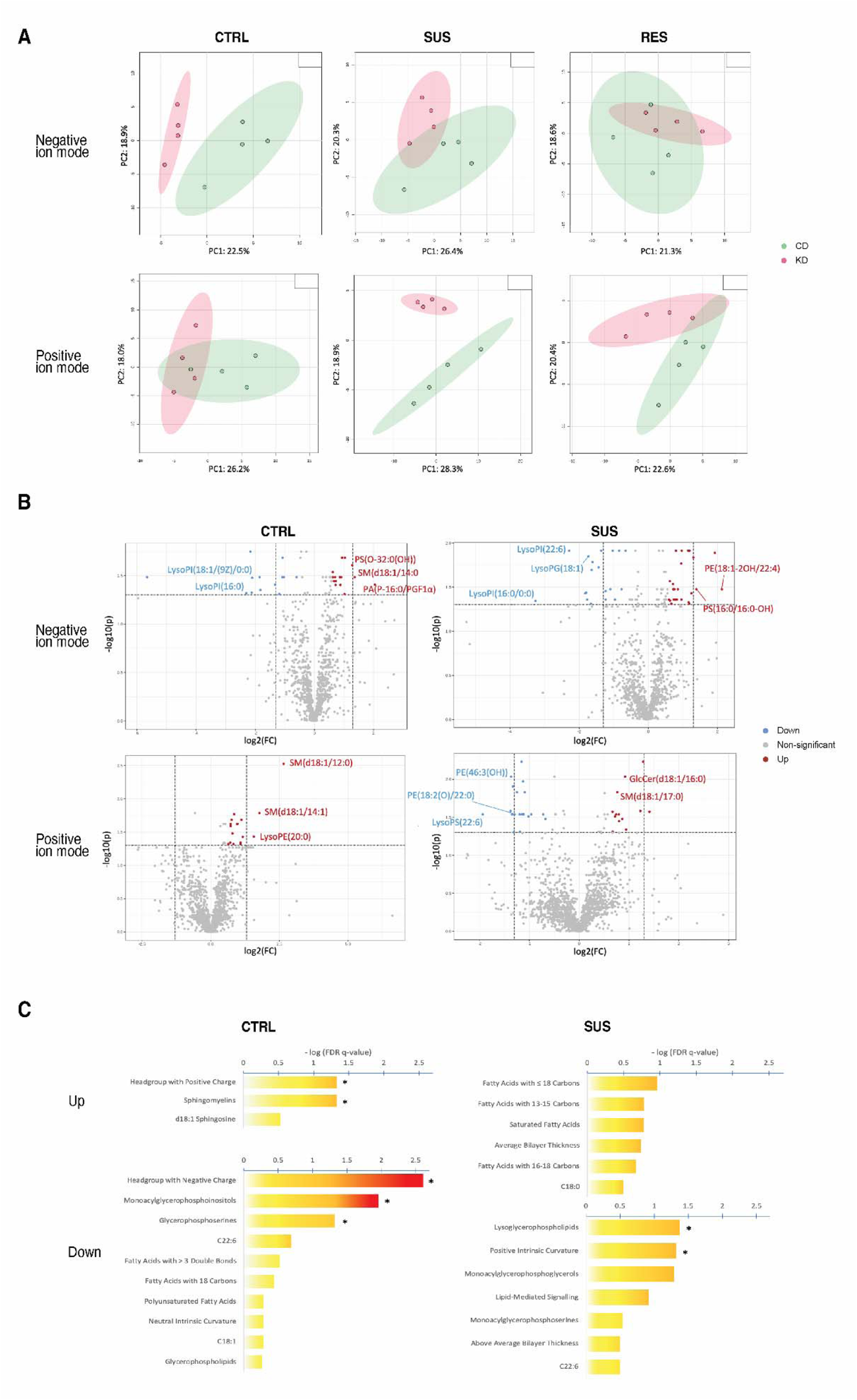
Ketogenic diet differentially alters hippocampal lipidomic profiles in non-stressed mice and mice exposed to repeated social defeat stress. PCA plots showing the separation in CTRL, SUS and RES groups, in positive and negative ion modes, respectively **(A)**. Volcano plots showing hippocampal differentially regulated lipids by a ketogenic diet (KD) *versus* control diet (CD) in the CTRL or SUS mice. Lipids indicated in red (up) or blue (down) are differentially regulated lipids, identified based on a fold change, FC >1.5 and a two-sample t-test p-value ≤0.05 that was considered significant **(B)**. Ontological analysis of differentially regulated lipids between KD and CD diets in CTRL and SUS mice. Lipids identified through two-sample t-test analyses as differentially regulated were annotated by matching to the human metabolome database (HMDB), followed by analysis in LION software in ranking mode. Lipid ontology terms that displayed differential regulation were considered to be significantly enriched (up) or underrepresented (down) are indicated by an asterisk (*) **(C)**. n = 4 mice/group. CD: control diet; KD: ketogenic diet; CTRL: control; SUS: susceptible; RES: resistant; GlcCer, glucosylceramide; PA, phosphatidic acid; PE, phosphatidylethanolamine; PGF1α, prostaglandin F1α; PI, phosphatidylinositol; PIP, phosphatidylinositol-monophosphate; PS, phosphatidylserine; SM, sphingomyelin.

Lipid ontology analyses using LION software (v.2020.07.14) ^85^ were performed to investigate the functional relevance of the DRLs identified (Fig. 6C, Supplementary Figure 1B, Supplementary Table 2). Among the lipids assigned with HMDB, 29 of 32 (91%) DRLs identified between diet groups in CTRL mice, 17 of 21 (81%) DRLs in RES mice, and 48 of 52 (92%) in SUS mice, were matched to the LION database. In CTRL mice, lipids with positively charged head groups (effect size (ES) = 0.615, q = 0.047) and ceramide phosphocholines (ES = 0.923, q = 0.047) were significantly enriched in the KD group, while lipids with negatively charged head groups (ES = −0.800, q = 0.002), monoacyl glycerophosphoinositols (ES = −1.00, q = 0.011) and glycerophosphoserines (ES = −0.840, q = 0.047) were significantly underrepresented. In SUS mice, lysoglycerophospholipids (ES = −0.615, q = 0.0422) and lipids with a positive intrinsic curvature (ES = −0.571, q = 0.048) were significantly underrepresented in the KD group. No significant ontological differences between the KD and CD groups were identified in RES mice.

## 4. Discussion

Our study investigated the effects of KD, in which ketone bodies instead of glucose are used as a source of energy, on stress resistance as well as microglial properties and the lipidomic profile in the ventral hippocampus. Mice following a KD did not show reduced weight compared to mice on a CD, an observation in line with those of another study in which 8–10-week-old male mice on a KD were exposed to chronic unpredictable stress ^4^. After 4 weeks, mice on a KD had elevated circulating levels of the ketone body BHB, confirming that these mice were deriving energy from the oxidation of ketone bodies instead of glucose.

Characterization of our model revealed that after 10 days of RSD, KD tended to increase the proportion of adult male mice classified as RES, defined as presenting social behavior similar to unstressed CTRLs during the SI test, compared to mice following CD. We observed an effect of psychosocial stress on blood CORT, with SUS mice showing increased levels compared to CTRL and RES animals. RSD also caused a transient elevation at Day 6 in blood levels of the inflammatory cytokines G-CSF, IL-6, IL-13 and IP-10 in SUS animals compared to CTRL. G-CSF was previously shown to have anti-inflammatory and neuroprotective properties in the brain ^96^. IL-6 is responsible for stimulating the production of C-reactive protein, a protein serving as a general marker of inflammation that correlates with depressive symptoms ^97^. IL-13 is produced by peripheral immune cells and microglia ^98,99^ and can exacerbate or resolve central nervous system inflammation, depending on the context ^99^. By contrast, IFN-g, a key mediator of microglial inflammatory response that was shown to drive depressive-like behavior in mice ^100,101^, presented no significant variation with the diet or stress paradigm. Nevertheless, we observed changes in IFN-g-related cytokines such as IP-10, which is increased in blood samples of patients with Alzheimer’s disease (AD) and in cerebrospinal fluid of patients with frontotemporal dementia encephalitis ^102,103^. A study showed that 10 days of RSD caused an increased number of peripheral neutrophils and monocytes in adult male mice ^104^. In our study, IFN-g levels were also reduced in the blood of mice following a KD.

The 10 days of RSD did not induce changes in microglial density and distribution in the CA1 *stratum radiatum*. Unchanged microglial density was previously reported in the same hippocampal region and layer in male mice following other protocols of stress such as chronic unpredictable mild stress ^62,105^. A limitation of the density analysis in our and previous studies might arise from the fact that different microglial states downregulating classical microglial markers might not be captured by Iba1 staining (combined with TMEM119 in this case) ^92^. However, our study revealed that microglia from KD-fed mice showed morphological changes at baseline, with larger soma and increased arborization area, within the CA1 *stratum radiatum*. It was previously observed that a KD on its own, at steady-state, can modify microglial morphology. In particular, microglia from the dorsal hippocampus of KD-fed adult male rats tended to have increased branching ^106^. In our study, microglia displayed increased soma and increased arborization without affecting the size ratio between soma and arborization with a KD, as seen with the morphological index in CTRL mice on both diets. Together, microglia of mice on a KD presented morphological adaptations associated to a tendency to have an increased prevalence of RES phenotype under this diet.

We also observed microglial morphological changes linked to the stress phenotypes, hinting to specific stress-induced microglial adaptations that could underlie these phenotypes. We observed an increase of microglial soma size in KD CTRL and CD SUS animals, compared to CD CTRL animals. In the same region, CA1 *stratum radiatum*, microglia with enlarged soma were previously described in an adult male mouse model of chronic unpredictable mild stress ^62^. Increased microglial soma area has been linked to elevated production of trophic factors and pro- or anti-inflammatory mediators ^91,105^, which could explain the observed soma increase associated to KD in the KD CTRL group and associated to stress in the CD SUS group. We further observed increased microglial arborization area at baseline in KD CTRL and in CD RES (CD CTRL *versus* CD RES) animals. This aligns with a surveillant function, considering that hyper-ramified microglia were associated with a stress resistant phenotype in the context of chronic unpredictable stress ^4^. A consequence of hyper-ramification is the higher number of functional contacts between microglia and the brain parenchyma ^107^. A recent study examining microglia in the CA1 region linked hyper-ramified microglia to stress resilience in a male mouse model of RSD ^108^. Similar observations were made in the habenula, a region involved in aversive negative behavior ^109^. When studying the effect of RSD, microglia in the habenula of SUS male mice showed increased cell body and decreased arborization volumes ^7^. In our study, no differences in microglial cell body and arborization area were linked to KD at steady-state, when comparing CD CTRL and CD RES mice. However, microglia from KD-fed *versus* CD-fed SUS mice were previously shown to display reduced cell body volume and increased arborization volume, highlighting the capacity of a KD to restore microglial morphology in a context of RSD ^7^.

Ultrastructural characterization of microglia by SEM allowed us to investigate at the nanoscale level possible changes in microglial intracellular organelles and contacts with specific elements of the parenchyma. SEM analysis showed that microglia of KD *versus* CD mice had a reduction of both empty phagosomes and tertiary lysosomes, hinting at possibly decreased contents that require degradation with the dietary change ^76,110^. Microglial cell bodies can establish functional membrane-to-membrane contacts with various types of elements although their specific functions remain to be more deeply understood ^94,111^. In our study, we observed microglial contacts with pre-synaptic axon terminals, dendritic spines and synaptic clefts, and with all categories combined. Ultrastructural analysis showed that microglia of SUS mice make fewer contacts with pre-synaptic axon terminals, post-synaptic dendritic spines and synaptic clefts compared to CTRL microglia. Additionally, we observed that microglia of SUS mice made overall less direct contacts with excitatory synapses. Given that further ultrastructural characterization of SUS mice did not reveal changes in microglial organelles involved in phagolysosomal activity, these reduced contacts were probably not associated to an active synaptic elimination at the time examined. Additional electrophysiological studies are warranted to determine the outcomes of these microglial contacts on synaptic density over the course of RSD.

We also found that microglia of mice following a KD had a reduced prevalence of cellular stress markers, seen as a reduced number of dilated Golgi apparatus and ER cisternae, as well as ER cisternae with dystrophy, and microglia with a lower percentage of Golgi apparatus displaying dilation. Elevations in reactive oxygen species (ROS), which are an important source of cellular stress, were shown to compromise the function of several cytoplasmic organelles ^77,112^. This could lead to impaired microglial function and a shift towards detrimental microglial states ^73^. The ketone body BHB can protect neurons from calcium-induced excitotoxicity and was shown to reduce reactive oxygen species (ROS) levels in neurons isolated from the rat brain ^113^. Increased ROS production in the brain was also linked to psychiatric diseases including mood disorders and major depression ^114^. Interestingly, one study showed that the behavioral changes associated with RSD require microglia and are driven by oxidative stress in adult male mice ^115^.

Microglial function is tightly regulated by mitochondrial dynamics ^116^. The structure of the mitochondrial network tightly influences microglial metabolism and their function, notably in response to pathological stimuli ^116,117^. Our analysis showed a reduction in the number of mitochondria in microglia from SUS mice on a CD diet, hinting that metabolic alterations in the CA1 region may contribute to the SUS phenotype under CD. We did not observe a different prevalence of dystrophic mitochondria in microglia from mice undergoing RSD, but an increased frequency of elongated mitochondria was found, suggesting mitochondrial adaptations. Mitochondria of different lengths were proposed to have different functions and energetic properties, including energy production efficiency ^118–120^. Further 3D analysis of the mitochondrial network would provide better understanding of the complex relationships between mitochondrial structure and function in microglia ^121^. Further analyses of functional mitochondrial output would help linking the ultrastructural modifications of mitochondria to their energetic proficiency among the different behavioral phenotypes.

Although glucose is the primary source of energy for the brain, microglial lipid metabolism is emerging as a key player in neuroimmune homeostasis ^122^. Microglia exist in a variety of homeostatic and inflammatory states, while recent evidence indicates that microglial uptake of lipids drives oxidative metabolism and an anti-inflammatory phenotype ^50,123^, and FAO-related genes are repressed in response to proinflammatory stimuli ^124^. Microglia take up free FAs through the scavenger receptor CD36 and import the cholesterol-containing apolipoproteins ApoE and ApoA1 through lipoprotein lipase (LPL) and the low-density lipoprotein receptor (LDL-R) ^122^. The uptake and subsequent degradation of microglial lipids is critical for the clearance of cellular debris, while deficiency in the triggering receptor expressed on myeloid cells 2 (TREM2), a key microglial sensor of apoptotic cells and extracellular debris, impairs microglial function in pathological states ^125^. As discussed below, our lipidomic data suggest an altered lipid profile in response to a KD that may reduce brain inflammation, promoting a homeostatic microglial state and supporting the possibility that microglia may underlie the proposed benefits of KD treatment for MDD.

Here, we identified profound changes in hippocampal lipid profiles in response to a KD, which differed among CTRL, SUS and RES mice. In general, a far greater number of lipids were differentially regulated by a KD in SUS mice than in CTRL or RES mice, potentially indicating that lipid metabolism in SUS mice varies in response to psychological stress, which may facilitate beneficial changes introduced by interventions such as a KD. In CTRL and RES mice, which may represent a more homeostatic state, a KD resulted in fewer changes to hippocampal lipid profile. It tended to broadly upregulate individual lipids, which suggests a general enrichment consistent with a lipid-rich diet. Perhaps surprisingly, a KD intervention did not significantly enrich any lipid ontology terms in RES mice, likely due to the low number of DRLs identified in these groups.

Lipid ontology analyses identified several terms significantly enriched by a KD across both CTRL and SUS mice. A KD treatment upregulated sphingomyelins, with SM(d18:1) species also trending upwards. These results are strikingly similar to the known effects of the antidepressants amitriptyline and fluoxetine, which were shown to induce sphingomyelin accumulation in an autophagy-dependent manner in cultured hippocampal neurons, thus promoting neurogenesis ^126^. As impaired neurogenesis is a major pathological mechanism in mood disorders, the ability of a KD to induce sphingomyelin accumulation may be highly desirable in a clinical context. Similarly chronic unpredictable stress leads to decreased hippocampal sphingomyelins in rats, which correlated negatively with blood CORT levels ^127^, supporting the idea that KD treatment could normalize stress-related alterations in lipid profiles. However, we did not observe consistent upregulation of sphingomyelins in SUS mice under KD, suggesting that a longer treatment period or combining KD treatment with conventional antidepressants may be necessary to increase sphingomyelin levels.

In both CTRL and SUS mice, KD treatment significantly altered lipid ontology terms related to glycerophospholipids, including downregulation of monoacylglycerophosphoinositols and glycerophosphoserines in CTRL mice and monoacylglycerophosphoglycerols in the SUS group. Glycerophospholipids are highly prevalent components of cell membranes, dictating membrane structure and function, while some less abundant lipids of this class, such as glycerophosphoinositols and glycerophosphoserines, are involved in cell-cell communication and intracellular signaling pathways ^128^. Moreover, the charged headgroups of phospholipids play a key role in determining membrane structure, which can in turn modify cellular function ^129,130^. For example, lipid headgroups with various charges differentially influence the activity of transmembrane receptors, such as the stress-related beta-adrenergic receptor, by modulating protein stability ^131^. Our lipid ontology results in CTRL mice showed significant upregulation of lipids with positively charged headgroups, which include phosphatidylethanolamines and phosphatidylcholines, and downregulation of those with negatively charged headgroups, such as PSs, in response to a KD. Thus, KD produces important changes in cell membrane structure, and by extension, may modify cellular function.

Specific lipids can also act as potent mediators of pro- or anti-inflammatory effects. Lysoglycerophospholipids are generally considered proinflammatory, promoting MIP-2 production and chemotaxis to apoptotic cells in macrophages ^132,133^. Lipidomic analyses have identified lysophosphatidylcholines (LPCs) and LPEs as significantly upregulated in rodent models of chronic restraint stress and serotonin deficiency ^134,135^, consistent with a role for inflammation and immune reactivity in depressive disorders. Meanwhile, KD treatment is widely reported to exert anti-inflammatory effects in rodent models of depression, potentially through its actions on microglia ^7,50^. Importantly, our results identified lysoglycerophospholipids as significantly downregulated by a KD in SUS mice, implying that a KD may attenuate inflammation in part through reducing proinflammatory lipid species such as lysoglycerophospholipids. It is worth noting that another study demonstrated increases in LPC upon treatment with conventional antidepressants ^136^. However, this study investigated lipidomic changes in healthy, rather than stressed mice. Indeed, our analyses of CTRL mice generally showed an upregulation of individual LPC species in response to KD, highlighting potentially similar mechanisms of KD and conventional antidepressant action that may differ between healthy and stressed subjects.

Notably, our results indicate that PSs were generally downregulated by KD treatment, with lipid ontology analyses showing a significant decrease in glycerophosphoserines in CTRL mice and a decreasing trend in monoacylglycerophosphoserines in SUS mice. PSs, normally localized to the internal leaflet of the plasma membrane, are externalized in apoptotic cells, marking the cell for phagocytosis by macrophages and microglia ^137^. The microglial receptor TREM2 is critical for this homeostatic process, as a loss of TREM2 impairs PS-mediated synaptic pruning of new hippocampal neurons during development and adult neurogenesis ^138,139^. Thus, increased brain PS content could indicate either increased neuronal cell death or defective microglial clearance of apoptotic neurons, both of which may contribute to pathology ^122^. The decrease in PSs observed in our study suggests either decreased neuronal death, notably in the context of adult neurogenesis, or improved microglial homeostatic function, consistent with the established effects of the KD in promoting anti-inflammatory microglial states ^50^.

As a limitation, this study was done only in male mice given that the RSD paradigm used relies on male-to-male aggressivity. Nevertheless, recent modifications of this paradigm can circumvent these behavioral limitations and have allowed other groups to characterize stress related changes in the female mouse brain. Future work using these models would be key for the study of sex differences in the outcomes of KD on stress related brain changes ^140,141^. While we focused on whole hippocampus for lipidomic analysis, RSD further increased the activity of the prefrontal cortex, bed nucleus of stria terminalis, and periaqueductal gray in mice ^63^. As a consequence, the results from this study should not be extrapolated to other brain regions which may differ in their susceptibilities to stress. It will be important in the future to ensure their further characterization, as behavior is the result of concerted changes among brain regions in constant communication. Furthermore, inflammatory responses are exacerbated with aging. A study showed that RSD caused a stronger inflammatory reaction in older *versus* younger mice. When analyzing the supernatants harvested from cultured splenocytes, the samples belonging to older mice contained higher IL-6 and TNF-α compared to samples from younger male mice ^142^. It would be interesting to study whether RSD in older mice would result in similar SUS to RES ratios and whether KD would still be able to promote a RES phenotype.

A KD has shown promising results as a potential beneficial dietary regime for many health complications including epilepsy and mood improvement ^5,8,11–13^. Nevertheless, low adherence rates are a limitation with this diet ^143^. Therefore, research that sheds light onto the mechanisms underlying the described beneficial effects could identify and isolate the responsible chemical compounds which could eliminate the requirement to adhere to this dietary regime. Exogenous supplementation of diets with ketone-derived products have shown promising results in preclinical and clinical studies. Supplementation with a ketone body-containing drink improved cognition in patients with mild cognitive impairment ^144^, while another study revealed that supplementation with ketogenic medium chain triglycerides increased energy metabolism in AD ^145^.

## 5. Conclusion

In conclusion, KD promoted resistance to psychological stress, by tending to increase the number of mice that did not show reduced sociability after 10 days of RSD. Our model allowed us to study specific changes linked to KD as well as study differences between the SUS and RES phenotypes under both diets. We further identified a transient elevation of circulating inflammatory cytokines, changes in microglial morphology, ultrastructural evidence of altered microglial direct contacts with synapses, cytoplasmic organelles, and cellular stress markers, as well as hippocampal lipidic expression due to stress and diet. Together, our results shed light onto the complex relationships between diet, immune system and stress resistance, contributing to a deeper understanding of the effects of stress and diet on the brain and behavior.

## 6. Acknowledgements

We thank all the personal of the animal facility of Centre de Recherche CHU de Quebec-Université Laval. We are grateful to the UVic Genome BC Proteomics Center for the lipidomic MS experiments. We also acknowledge with respect the lək□□əŋən peoples on whose traditional territory the University of Victoria stands and the Songhees, Esquimalt and WlJSÁNEĆ peoples whose historical relationships with the land continue to this day.

We further acknowledge all the funding agencies making this work possible. F.G.I. was supported by a full doctoral scholarship from the Mexican Council of Science and Technology (CONACYT). T.H. was supported by a Natural Sciences and Engineering Research Council of Canada (NSERC) Undergraduate Student Research Award. M.C. and K.P. were supported by Fonds de Recherche du Québec – Santé (FRQS) Doctoral Training Awards. K.P. was also supported by a Centre de Thématique de Recherche en Neurosciences scholarship. M.E.T. is a Tier II Canada Research Chair in *Neurobiology of Aging and Cognition*. J.D. was funded by FRQS Research Scholar Junior 1 #298905 and a National Institutes of Health (NIH) grant #1R21MH119561-01A1. M.L. was supported by the Academy of Finland grant #318857. This research was also funded by a NSERC Discovery grant awarded to M.E.T. (RGPIN-2014-05308) and the ERA-NET Co-Fund project JTC2017, MicroSynDep Consortium, to J.D., M.L. and M.E.T.

## 7. Author contributions

K.S., N.V., F.G.I. and M.E.T. conceived the project and designed the experiments. Animal handling and diet control were done by K.S. and N.V. with the help of K.B. Behavioral experiments were done by K.S. N.V. and K.B. Tissue processing was performed by K.S., N.V., K.B., and F.G.I. The immunofluorescence staining, epifluorescence and confocal imaging, as well as analysis were done by C.M., K.P., and F.G.I. Electron microscopy image acquisition and analysis were done by F.G.I. In addition, T.H., M.C. and M.L. were in charge of the lipidomic sample processing and bioinformatic analyses. F.G.I. and T.H. prepared the manuscript, as well as figures and tables, under the supervision of M.E.T. Proof reading and approval of the final manuscript was done by F.G.I., T.H., J.D., M.L. and M.E.T.

## 8. Declaration of Competing Interest

The authors declare no competing interest.

## 9. Data Availability

The raw data supporting the findings of this study are available from the corresponding author upon reasonable request.

## Abbreviations

+: positive
AD: Alzheimer’s disease
ANOVA: analysis of variance
BB: blocking buffer
BHB: β-hydroxybutyrate
BV: blood vessel
CA1: *cornu ammonis*
CD: control diet
CID: collision induced dissociation
CORT: corticosterone
CTRL: control
CZ: corner zone
DAB: 3,3’ diaminobenzidine
DRLs: differentially-regulated lipids
ER: endoplasmic reticulum
ES: effect size
FC: fold change
FDS: false discovery range
FWHM: full width at half maximum
G-CSF: granulocyte colony stimulating factor
GlcCer: glucosylceramide
GM-CSF: granulocyte macrophage colony stimulating factor
HMDB: human metabolome data base
Iba1: ionized calcium-binding adapter molecule 1
IFN-□: interferon gamma
IL: interleukin
iNOS: inducible nitric oxide synthase
IP-10: C-X-C motif chemokine ligand 10
IQR: interquartile range
IZ: interaction zone
KC: keratinocytes-derived chemokine
KD: ketogenic diet
LC/MS: Liquid Chromatography/Mass Spectrometry
LDL-R: low-density lipoprotein receptor
LIF: leukemia inhibitory factor
LIX: lipopolysaccharide-induced CXC chemokine
LPCs: lysophosphatidylcholines
LPEs: lysophosphatidylethanolamines
LPL: lipoprotein lipase
LPS: lipopolysaccharide
M-CSF: macrophage colony stimulating factor
m/z: mass to charge ratio
MCP-1: monocyte chemoattractant protein-1
MDD: major depressive disorder
MIG: monokine induced by interferon-gamma
MIP-1α: macrophage inflammatory protein 1-alpha
MIP-1β: macrophage inflammatory protein 1-beta
MIP-2: macrophage inflammatory protein 2
n: sample size
NF-kb: nuclear factor kappa b
NLPR3: nucleotide-binding and oligomerization domain-like receptors pyrin domain-containing protein 3
NND: nearest neighbor distance
PA: phosphatidic acid
PB: phosphate buffer
PBS: phosphate-buffered saline
PBST: PBS containing Triton X-100
PC: principal component
PE: phosphatidylethanolamine
PFA: paraformaldehyde
PGF1α: prostaglandin F1 alpha
PI: phosphatidylinositol
PIP: phosphatidylinositol-monophosphate
PSs: phosphatidylserines
QC: quality control
RANTES: C–C chemokine ligand 5
RES: resistant
ROS: reactive oxygen species
RSD: repeated social defeat
RSDs: relative standard deviations
RT: room temperature
SEM: scanning electron microscope
SI: social interaction
SM: sphingomyelin
SUS: susceptible
TB: Tris-buffer
TBS: Tris-buffered saline
TMEM119: transmembrane protein 119
TNF-α: tumor necrosis factor alpha
TREM2: triggering receptor expressed on myeloid cells 2
VGEF: vascular endothelial growth factor

## Notes

### Competing Interest Statement

The authors have declared no competing interest.

## References

1. Brietzke E, Mansur RB, Subramaniapillai M, et al. Ketogenic diet as a metabolic therapy for mood disorders: Evidence and developments. Neuroscience & Biobehavioral Reviews. 2018;94:11–16. doi:10.1016/j.neubiorev.2018.07.020

2. Opie RS, O’Neil A, Jacka FN, Pizzinga J, Itsiopoulos C. A modified Mediterranean dietary intervention for adults with major depression: Dietary protocol and feasibility data from the SMILES trial. Nutritional Neuroscience. 2018;21(7):487–501. doi:10.1080/1028415X.2017.1312841

3. Igwe O, Sone M, Matveychuk D, Baker GB, Dursun SM. A review of effects of calorie restriction and fasting with potential relevance to depression. Progress in Neuro-Psychopharmacology and Biological Psychiatry. 2021;111:110206. doi:10.1016/j.pnpbp.2020.110206

4. Huang C, Wang P, Xu X, et al. The ketone body metabolite β-hydroxybutyrate induces an antidepression-associated ramification of microglia via HDACs inhibition-triggered Akt-small RhoGTPase activation. Glia. 2018;66(2):256–278. 10.1002/glia.23241

5. Ricci A, Idzikowski MA, Soares CN, Brietzke E. Exploring the mechanisms of action of the antidepressant effect of the ketogenic diet. Reviews in the Neurosciences. 2020;31(6):637–648. doi:10.1515/revneuro-2019-0073

6. MartinLMcGill KJ, Jackson CF, Bresnahan R, Levy RG, Cooper PN. Ketogenic diets for drugLresistant epilepsy. Cochrane Database Syst Rev. 2018;2018(11):CD001903. doi:10.1002/14651858.CD001903.pub4

7. Guan YF, Huang GB, Xu MD, et al. Anti-depression effects of ketogenic diet are mediated via the restoration of microglial activation and neuronal excitability in the lateral habenula. Brain, Behavior, and Immunity. Published online May 12, 2020. doi:10.1016/j.bbi.2020.05.032

8. Włodarczyk A, Cubała WJ, Stawicki M. Ketogenic diet for depression: A potential dietary regimen to maintain euthymia? Progress in Neuro-Psychopharmacology and Biological Psychiatry. 2021;109:110257. doi:10.1016/j.pnpbp.2021.110257

9. Murphy P, Likhodii S, Nylen K, Burnham WM. The antidepressant properties of the ketogenic diet. Biological Psychiatry. 2004;56(12):981–983. doi:10.1016/j.biopsych.2004.09.019

10. Nei M, Ngo L, Sirven JI, Sperling MR. Ketogenic diet in adolescents and adults with epilepsy. Seizure. 2014;23(6):439–442. doi:10.1016/j.seizure.2014.02.015

11. Ashton JS, Roberts JW, Wakefield CJ, et al. The effects of medium chain triglyceride (MCT) supplementation using a C8:C10 ratio of 30:70 on cognitive performance in healthy young adults. Physiology & Behavior. 2021;229:113252. doi:10.1016/j.physbeh.2020.113252

12. Carneiro L, Pellerin L. Nutritional Impact on Metabolic Homeostasis and Brain Health. Front Neurosci. 2022;15:767405. doi:10.3389/fnins.2021.767405

13. Jiwani R, Robbins R, Neri A, Renero J, Lopez E, Serra MC. Effect of Dietary Intake Through Whole Foods on Cognitive Function: Review of Randomized Controlled Trials. Curr Nutr Rep. 2022;11(2):146–160. doi:10.1007/s13668-022-00412-5

14. Mergenthaler P, Lindauer U, Dienel GA, Meisel A. Sugar for the brain: the role of glucose in physiological and pathological brain function. Trends Neurosci. 2013;36(10):587–597. doi:10.1016/j.tins.2013.07.001

15. Boison D. New insights into the mechanisms of the ketogenic diet. Curr Opin Neurol. 2017;30(2):187–192. doi:10.1097/WCO.0000000000000432

16. de Lima PA, de Brito Sampaio LP, Damasceno NRT. Neurobiochemical mechanisms of a ketogenic diet in refractory epilepsy. Clinics (Sao Paulo). 2014;69(10):699–705. doi:10.6061/clinics/2014(10)09

17. IJff DM, Postulart D, Lambrechts DAJE, et al. Cognitive and behavioral impact of the ketogenic diet in children and adolescents with refractory epilepsy: A randomized controlled trial. Epilepsy & Behavior. 2016;60:153–157. doi:10.1016/j.yebeh.2016.04.033

18. Clark PJ, Brzezinska WJ, Puchalski EK, Krone DA, Rhodes JS. Functional Analysis of Neurovascular Adaptations to Exercise in the Dentate Gyrus of Young Adult Mice Associated With Cognitive Gain. Hippocampus. 2009;19(10):937–950. doi:10.1002/hipo.20543

19. Brown ES, Hughes CW, McColl R, Peshock R, King KS, Rush AJ. Association of Depressive Symptoms with Hippocampal Volume in 1936 Adults. Neuropsychopharmacology. 2014;39(3):770–779. doi:10.1038/npp.2013.271

20. Parfitt GM, Nguyen R, Bang JY, et al. Bidirectional Control of Anxiety-Related Behaviors in Mice: Role of Inputs Arising from the Ventral Hippocampus to the Lateral Septum and Medial Prefrontal Cortex. Neuropsychopharmacology. 2017;42(8):1715–1728. doi:10.1038/npp.2017.56

21. Vyas A, Mitra R, Shankaranarayana Rao BS, Chattarji S. Chronic Stress Induces Contrasting Patterns of Dendritic Remodeling in Hippocampal and Amygdaloid Neurons. J Neurosci. 2002;22(15):6810–6818. doi:10.1523/JNEUROSCI.22-15-06810.2002

22. Schoenfeld TJ, McCausland HC, Morris HD, Padmanaban V, Cameron HA. Stress and loss of adult neurogenesis differentially reduce hippocampal volume. Biol Psychiatry. 2017;82(12):914–923. doi:10.1016/j.biopsych.2017.05.013

23. Campbell S, MacQueen G. The role of the hippocampus in the pathophysiology of major depression. J Psychiatry Neurosci. 2004;29(6):417–426.

24. Videbech P, Ravnkilde B. Hippocampal Volume and Depression: A Meta-Analysis of MRI Studies. AJP. 2004;161(11):1957–1966. doi:10.1176/appi.ajp.161.11.1957

25. Woodburn SC, Bollinger JL, Wohleb ES. The semantics of microglia activation: neuroinflammation, homeostasis, and stress. J Neuroinflammation. 2021;18(1):1–16. doi:10.1186/s12974-021-02309-6

26. Magarinos AM, McEwen BS. Stress-induced atrophy of apical dendrites of hippocampal CA3c neurons: Comparison of stressors. Neuroscience. 1995;69(1):83–88. doi:10.1016/0306-4522(95)00256-I

27. Frodl T, Meisenzahl EM, Zetzsche T, et al. Hippocampal changes in patients with a first episode of major depression. Am J Psychiatry. 2002;159(7):1112–1118. doi:10.1176/appi.ajp.159.7.1112

28. Holmes SE, Scheinost D, Finnema SJ, et al. Lower synaptic density is associated with depression severity and network alterations. Nat Commun. 2019;10(1):1529. doi:10.1038/s41467-019-09562-7

29. Gold PW. The organization of the stress system and its dysregulation in depressive illness. Mol Psychiatry. 2015;20(1):32–47. doi:10.1038/mp.2014.163

30. Saveanu RV, Nemeroff CB. Etiology of Depression: Genetic and Environmental Factors. Psychiatric Clinics of North America. 2012;35(1):51–71. doi:10.1016/j.psc.2011.12.001

31. Beurel E, Toups M, Nemeroff CB. The Bidirectional Relationship of Depression and Inflammation: Double Trouble. Neuron. 2020;107(2):234–256. doi:10.1016/j.neuron.2020.06.002

32. Casaril AM, Dantzer R, Bas-Orth C. Neuronal Mitochondrial Dysfunction and Bioenergetic Failure in Inflammation-Associated Depression. Front Neurosci. 2021;15:725547. doi:10.3389/fnins.2021.725547

33. Krishnan V, Han MH, Graham DL, et al. Molecular adaptations underlying susceptibility and resistance to social defeat in brain reward regions. Cell. 2007;131(2):391–404. doi:10.1016/j.cell.2007.09.018

34. Miller AH, Raison CL. The role of inflammation in depression: from evolutionary imperative to modern treatment target. Nat Rev Immunol. 2016;16(1):22–34. doi:10.1038/nri.2015.5

35. Kitaoka S. Inflammation in the brain and periphery found in animal models of depression and its behavioral relevance. Journal of Pharmacological Sciences. 2022;148(2):262–266. doi:10.1016/j.jphs.2021.12.005

36. Koo JW, Wohleb ES. How stress shapes neuroimmune function: implications for the neurobiology of psychiatric disorders. Biol Psychiatry. 2021;90(2):74–84. doi:10.1016/j.biopsych.2020.11.007

37. Bonaccorso S, Marino V, Puzella A, et al. Increased Depressive Ratings in Patients With Hepatitis C Receiving Interferon-α–Based Immunotherapy Are Related to Interferon-α– Induced Changes in the Serotonergic System. Journal of Clinical Psychopharmacology. 2002;22(1):86.

38. Capuron L, Castanon N. Role of Inflammation in the Development of Neuropsychiatric Symptom Domains: Evidence and Mechanisms. In: Dantzer R, Capuron L, eds. Inflammation-Associated Depression: Evidence, Mechanisms and Implications. Current Topics in Behavioral Neurosciences. Springer International Publishing; 2017:31–44. doi:10.1007/7854_2016_14

39. Walker FR, Beynon SB, Jones KA, et al. Dynamic structural remodelling of microglia in health and disease: A review of the models, the signals and the mechanisms. Brain, Behavior, and Immunity. 2014;37:1–14. doi:10.1016/j.bbi.2013.12.010

40. Eisenberger NI, Berkman ET, Inagaki TK, Rameson LT, Mashal NM, Irwin MR. Inflammation-Induced Anhedonia: Endotoxin Reduces Ventral Striatum Responses to Reward. Biol Psychiatry. 2010;68(8):748–754. doi:10.1016/j.biopsych.2010.06.010

41. Harrison NA, Brydon L, Walker C, Gray MA, Steptoe A, Critchley HD. Inflammation Causes Mood Changes Through Alterations in Subgenual Cingulate Activity and Mesolimbic Connectivity. Biol Psychiatry. 2009;66(5):407–414. doi:10.1016/j.biopsych.2009.03.015

42. Allison DJ, Sharma B, Timmons BW. The efficacy of anti-inflammatory treatment interventions on depression in individuals with major depressive disorder and high levels of inflammation: A systematic review of randomized clinical trials. Physiology & Behavior. 2019;207:104–112. doi:10.1016/j.physbeh.2019.05.006

43. Dantzer R, Cohen S, Russo SJ, Dinan TG. Resilience and immunity. Brain Behav Immun. 2018;74:28–42. doi:10.1016/j.bbi.2018.08.010

44. Tolkien K, Bradburn S, Murgatroyd C. An anti-inflammatory diet as a potential intervention for depressive disorders: A systematic review and meta-analysis. Clinical Nutrition. 2019;38(5):2045–2052. doi:10.1016/j.clnu.2018.11.007

45. Bohlen CJ, Friedman BA, Dejanovic B, Sheng M. Microglia in Brain Development, Homeostasis, and Neurodegeneration. Annu Rev Genet. 2019;53:263–288. doi:10.1146/annurev-genet-112618-043515

46. Carrier M, Šimončičová E, St-Pierre MK, McKee C, Tremblay MÈ. Psychological Stress as a Risk Factor for Accelerated Cellular Aging and Cognitive Decline: The Involvement of Microglia-Neuron Crosstalk. Front Mol Neurosci. 2021;14:749737. doi:10.3389/fnmol.2021.749737

47. Nimmerjahn A, Kirchhoff F, Helmchen F. Resting Microglial Cells Are Highly Dynamic Surveillants of Brain Parenchyma in Vivo. Science. 2005;308(5726):1314–1318. doi:10.1126/science.1110647

48. Kettenmann H, Hanisch UK, Noda M, Verkhratsky A. Physiology of Microglia. Physiological Reviews. 2011;91(2):461–553. doi:10.1152/physrev.00011.2010

49. Enache D, Pariante CM, Mondelli V. Markers of central inflammation in major depressive disorder: A systematic review and meta-analysis of studies examining cerebrospinal fluid, positron emission tomography and post-mortem brain tissue. Brain, Behavior, and Immunity. 2019;81:24–40. doi:10.1016/j.bbi.2019.06.015

50. Morris G, Puri BK, Maes M, Olive L, Berk M, Carvalho AF. The role of microglia in neuroprogressive disorders: mechanisms and possible neurotherapeutic effects of induced ketosis. Progress in Neuro-Psychopharmacology and Biological Psychiatry. 2020;99:109858. doi:10.1016/j.pnpbp.2020.109858

51. Zhang L, Zhang J, You Z. Switching of the Microglial Activation Phenotype Is a Possible Treatment for Depression Disorder. Front Cell Neurosci. 2018;12. doi:10.3389/fncel.2018.00306

52. Wohleb ES, McKim DB, Sheridan JF, Godbout JP. Monocyte trafficking to the brain with stress and inflammation: a novel axis of immune-to-brain communication that influences mood and behavior. Front Neurosci. 2015;8. doi:10.3389/fnins.2014.00447

53. Setiawan E, Wilson AA, Mizrahi R, et al. Increased Translocator Protein Distribution Volume, A Marker of Neuroinflammation, in the Brain During Major Depressive Episodes. JAMA Psychiatry. 2015;72(3):268–275. doi:10.1001/jamapsychiatry.2014.2427

54. Franklin TC, Wohleb ES, Zhang Y, Fogaça M, Hare B, Duman RS. Persistent increase in microglial RAGE contributes to chronic stress Induced priming of depressive-like behavior. Biol Psychiatry. 2018;83(1):50–60. doi:10.1016/j.biopsych.2017.06.034

55. Stratoulias V, Venero JL, Tremblay M, Joseph B. Microglial subtypes: diversity within the microglial community. EMBO J. 2019;38(17):e101997. doi:10.15252/embj.2019101997

56. Troubat R, Barone P, Leman S, et al. Neuroinflammation and depression: A review. European Journal of Neuroscience. 2021;53(1):151–171. doi:10.1111/ejn.14720

57. Toenders YJ, Laskaris L, Davey CG, et al. Inflammation and depression in young people: a systematic review and proposed inflammatory pathways. Mol Psychiatry. 2022;27(1):315–327. doi:10.1038/s41380-021-01306-8

58. Dupuis N, Curatolo N, Benoist JF, Auvin S. Ketogenic diet exhibits anti-inflammatory properties. Epilepsia. 2015;56(7):e95–98. doi:10.1111/epi.13038

59. Youm YH, Nguyen KY, Grant RW, et al. Ketone body β-hydroxybutyrate blocks the NLRP3 inflammasome-mediated inflammatory disease. Nat Med. 2015;21(3):263–269. doi:10.1038/nm.3804

60. Iwata M, Ota KT, Li XY, et al. Psychological Stress Activates the Inflammasome via Release of Adenosine Triphosphate and Stimulation of the Purinergic Type 2X7 Receptor. Biological Psychiatry. 2016;80(1):12–22. doi:10.1016/j.biopsych.2015.11.026

61. Rahimian R, Belliveau C, Chen R, Mechawar N. Microglial Inflammatory-Metabolic Pathways and Their Potential Therapeutic Implication in Major Depressive Disorder. Front Psychiatry. 2022;13:871997. doi:10.3389/fpsyt.2022.871997

62. Milior G, Lecours C, Samson L, et al. Fractalkine receptor deficiency impairs microglial and neuronal responsiveness to chronic stress. *Brain*, Behavior, and Immunity. 2016;55:114–125. doi:10.1016/j.bbi.2015.07.024

63. Laine MA, Sokolowska E, Dudek M, Callan SA, Hyytiä P, Hovatta I. Brain activation induced by chronic psychosocial stress in mice. Sci Rep. 2017;7:15061. doi:10.1038/s41598-017-15422-5

64. Bannerman DM, Grubb M, Deacon RMJ, Yee BK, Feldon J, Rawlins JNP. Ventral hippocampal lesions affect anxiety but not spatial learning. Behavioural Brain Research. 2003;139(1):197–213. doi:10.1016/S0166-4328(02)00268-1

65. Golden SA, Covington HE, Berton O, Russo SJ. A standardized protocol for repeated social defeat stress in mice. Nature Protocols. 2011;6(8):1183–1191. doi:10.1038/nprot.2011.361

66. Henry MS, Bisht K, Vernoux N, et al. Delta Opioid Receptor Signaling Promotes Resilience to Stress Under the Repeated Social Defeat Paradigm in Mice. Front Mol Neurosci. 2018;11. doi:10.3389/fnmol.2018.00100

67. Berton O, McClung CA, DiLeone RJ, et al. Essential Role of BDNF in the Mesolimbic Dopamine Pathway in Social Defeat Stress. Science. 2006;311(5762):864–868. doi:10.1126/science.1120972

68. Menard C, Pfau ML, Hodes GE, et al. Social stress induces neurovascular pathology promoting depression. Nat Neurosci. 2017;20(12):1752–1760. doi:10.1038/s41593-017-0010-3

69. Peng J, Liu Y, Umpierre AD, et al. Microglial P2Y12 receptor regulates ventral hippocampal CA1 neuronal excitability and innate fear in mice. Molecular Brain. 2019;12(1):71. doi:10.1186/s13041-019-0492-x

70. Bennett ML, Bennett FC, Liddelow SA, et al. New tools for studying microglia in the mouse and human CNS. Proc Natl Acad Sci U S A. 2016;113(12):E1738–E1746. doi:10.1073/pnas.1525528113

71. Ibanez FG, Picard K, Bordeleau M, Sharma K, Bisht K, Tremblay MÈ. Immunofluorescence Staining Using IBA1 and TMEM119 for Microglial Density, Morphology and Peripheral Myeloid Cell Infiltration Analysis in Mouse Brain. JoVE (Journal of Visualized Experiments). 2019;(152):e60510. doi:10.3791/60510

72. Alan Peters, Sanford Palay, Webster Henry. The Fine Structure of the Nervous System: Neurons and Their Supporting Cells, 3rd Edn. New York, NY: Oxford University Press.; 1990. Accessed November 25, 2021. http://onlinelibrary.wiley.com/doi/abs/10.1002/ana.410040660

73. Nahirney PC, Tremblay ME. Brain Ultrastructure: Putting the Pieces Together. Frontiers in Cell and Developmental Biology. 2021;9:187. doi:10.3389/fcell.2021.629503

74. Tremblay MÈ, Lowery RL, Majewska AK. Microglial Interactions with Synapses Are Modulated by Visual Experience. PLoS Biol. 2010;8(11). doi:10.1371/journal.pbio.1000527

75. St-Pierre MK, Carrier M, Lau V, Tremblay MÈ. Investigating Microglial Ultrastructural Alterations and Intimate Relationships with Neuronal Stress, Dystrophy, and Degeneration in Mouse Models of Alzheimer’s Disease. In: Jahani-Asl A, ed. Neuronal Cell Death: Methods and Protocols. Methods in Molecular Biology. Springer US; 2022:29–58. doi:10.1007/978-1-0716-2409-8_3

76. El Hajj H, Savage JC, Bisht K, et al. Ultrastructural evidence of microglial heterogeneity in Alzheimer’s disease amyloid pathology. J Neuroinflammation. 2019;16. doi:10.1186/s12974-019-1473-9

77. Chavez-Valdez R, Flock DL, Martin LJ, Northington FJ. Endoplasmic Reticulum pathology and stress response in neurons precede programmed necrosis after neonatal hypoxia-ischemia. Int J Dev Neurosci. 2016;48:58–70. doi:10.1016/j.ijdevneu.2015.11.007

78. Savage JC, St-Pierre MK, Carrier M, et al. Microglial physiological properties and interactions with synapses are altered at presymptomatic stages in a mouse model of Huntington’s disease pathology. J Neuroinflammation. 2020;17(1):98. doi:10.1186/s12974-020-01782-9

79. Lecours C, St-Pierre MK, Picard K, et al. Levodopa partially rescues microglial numerical, morphological, and phagolysosomal alterations in a monkey model of Parkinson’s disease. Brain, Behavior, and Immunity. 2020;90:81–96. doi:10.1016/j.bbi.2020.07.044

80. Benton HP, Want EJ, Ebbels TMD. Correction of mass calibration gaps in liquid chromatography–mass spectrometry metabolomics data. Bioinformatics. 2010;26(19):2488–2489. doi:10.1093/bioinformatics/btq441

81. Smith CA, Want EJ, O’Maille G, Abagyan R, Siuzdak G. XCMS:L Processing Mass Spectrometry Data for Metabolite Profiling Using Nonlinear Peak Alignment, Matching, and Identification. Anal Chem. 2006;78(3):779–787. doi:10.1021/ac051437y

82. Tautenhahn R, Böttcher C, Neumann S. Highly sensitive feature detection for high resolution LC/MS. BMC Bioinformatics. 2008;9(1):504. doi:10.1186/1471-2105-9-504

83. Kind T, Fiehn O. Seven Golden Rules for heuristic filtering of molecular formulas obtained by accurate mass spectrometry. BMC Bioinformatics. 2007;8:105. doi:10.1186/1471-2105-8-105

84. Pang Z, Zhou G, Ewald J, et al. Using MetaboAnalyst 5.0 for LC–HRMS spectra processing, multi-omics integration and covariate adjustment of global metabolomics data. Nat Protoc. 2022;17(8):1735–1761. doi:10.1038/s41596-022-00710-w

85. Molenaar MR, Jeucken A, Wassenaar TA, van de Lest CHA, Brouwers JF, Helms JB. LION/web: a web-based ontology enrichment tool for lipidomic data analysis. Gigascience. 2019;8(6):giz061. doi:10.1093/gigascience/giz061

86. Hui CW, St-Pierre A, El Hajj H, et al. Prenatal Immune Challenge in Mice Leads to Partly Sex-Dependent Behavioral, Microglial, and Molecular Abnormalities Associated with Schizophrenia. Front Mol Neurosci. 2018;11:13. doi:10.3389/fnmol.2018.00013

87. Li Q, Cheng Z, Zhou L, et al. Developmental Heterogeneity of Microglia and Brain Myeloid Cells Revealed by Deep Single-Cell RNA Sequencing. Neuron. 2019;101(2):207–223.e10. doi:10.1016/j.neuron.2018.12.006

88. Cunningham C, Wilcockson DC, Campion S, Lunnon K, Perry VH. Central and Systemic Endotoxin Challenges Exacerbate the Local Inflammatory Response and Increase Neuronal Death during Chronic Neurodegeneration. J Neurosci. 2005;25(40):9275–9284. doi:10.1523/JNEUROSCI.2614-05.2005

89. Picard K, Bisht K, Poggini S, et al. Microglial-glucocorticoid receptor depletion alters the response of hippocampal microglia and neurons in a chronic unpredictable mild stress paradigm in female mice. *Brain*, Behavior, and Immunity. 2021;97:423–439. doi:10.1016/j.bbi.2021.07.022

90. Tay TL, Savage JC, Hui CW, Bisht K, Tremblay M. Microglia across the lifespan: from origin to function in brain development, plasticity and cognition. J Physiol. 2017;595(6):1929–1945. doi:10.1113/JP272134

91. Savage JC, St-Pierre MK, Hui CW, Tremblay ME. Microglial Ultrastructure in the Hippocampus of a Lipopolysaccharide-Induced Sickness Mouse Model. Front Neurosci. 2019;13:1340. doi:10.3389/fnins.2019.01340

92. Paolicelli RC, Sierra A, Stevens B, et al. Microglia states and nomenclature: A field at its crossroads. Neuron. 2022;110(21):3458–3483. doi:10.1016/j.neuron.2022.10.020

93. Gonçalves de Andrade E, González Ibáñez F, Tremblay MÈ. Microglia as a Hub for Suicide Neuropathology: Future Investigation and Prevention Targets. Front Cell Neurosci. 2022;16:839396. doi:10.3389/fncel.2022.839396

94. Cserép C, Pósfai B, Dénes Á. Shaping Neuronal Fate: Functional Heterogeneity of Direct Microglia-Neuron Interactions. Neuron. 2021;109(2):222–240. doi:10.1016/j.neuron.2020.11.007

95. Umpierre AD, Bystrom LL, Ying Y, Liu YU, Worrell G, Wu LJ. Microglial calcium signaling is attuned to neuronal activity in awake mice. Bergles DE, Aldrich RW, eds. eLife. 2020;9:e56502. doi:10.7554/eLife.56502

96. Rahi V, Jamwal S, Kumar P. Neuroprotection through G-CSF: recent advances and future viewpoints. Pharmacol Rep. 2021;73(2):372–385. doi:10.1007/s43440-020-00201-3

97. Kappelmann N, Arloth J, Georgakis MK, et al. Dissecting the Association Between Inflammation, Metabolic Dysregulation, and Specific Depressive Symptoms: A Genetic Correlation and 2-Sample Mendelian Randomization Study. JAMA Psychiatry. 2021;78(2):161–170. doi:10.1001/jamapsychiatry.2020.3436

98. Ellis SL, Gysbers V, Manders PM, et al. The Cell-Specific Induction of CXC Chemokine Ligand 9 Mediated by IFN-γ in Microglia of the Central Nervous System Is Determined by the Myeloid Transcription Factor PU.1. J Immunol. 2010;185(3):1864–1877. doi:10.4049/jimmunol.1000900

99. Miao W, Zhao Y, Huang Y, et al. IL-13 Ameliorates Neuroinflammation and Promotes Functional Recovery after Traumatic Brain Injury. J Immunol. 2020;204(6):1486–1498. doi:10.4049/jimmunol.1900909

100. Zhang J, Rong P, Zhang L, et al. IL4-driven microglia modulate stress resilience through BDNF-dependent neurogenesis. Sci Adv. 2021;7(12):eabb9888. doi:10.1126/sciadv.abb9888

101. Kann O, Almouhanna F, Chausse B. Interferon γ: a master cytokine in microglia-mediated neural network dysfunction and neurodegeneration. Trends in Neurosciences. 2022;45(12):913–927. doi:10.1016/j.tins.2022.10.007

102. Curtin NM, Boyle NT, Mills KHG, Connor TJ. Psychological stress suppresses innate IFN-γ production via glucocorticoid receptor activation: Reversal by the anxiolytic chlordiazepoxide. *Brain*, Behavior, and Immunity. 2009;23(4):535–547. doi:10.1016/j.bbi.2009.02.003

103. Galimberti D, Bonsi R, Fenoglio C, et al. Inflammatory molecules in Frontotemporal Dementia: Cerebrospinal fluid signature of progranulin mutation carriers. *Brain*, Behavior, and Immunity. 2015;49:182–187. doi:10.1016/j.bbi.2015.05.006

104. Ishikawa Y, Kitaoka S, Kawano Y, et al. Repeated social defeat stress induces neutrophil mobilization in mice: maintenance after cessation of stress and strain-dependent difference in response. British Journal of Pharmacology. 2021;178(4):827–844. doi:10.1111/bph.15203

105. Alboni S, Poggini S, Garofalo S, et al. Fluoxetine treatment affects the inflammatory response and microglial function according to the quality of the living environment. Brain, Behavior, and Immunity. 2016;58:261–271. doi:10.1016/j.bbi.2016.07.155

106. Gzielo K, Soltys Z, Rajfur Z, Setkowicz ZK. The Impact of the Ketogenic Diet on Glial Cells Morphology. A Quantitative Morphological Analysis. Neuroscience. 2019;413:239–251. doi:10.1016/j.neuroscience.2019.06.009

107. Cserép C, Pósfai B, Lénárt N, et al. Microglia monitor and protect neuronal function via specialized somatic purinergic junctions. Science. Published online December 12, 2019. doi:10.1126/science.aax6752

108. Fujikawa R, Jinno S. Identification of hyper-ramified microglia in the CA1 region of the mouse hippocampus potentially associated with stress resilience. European Journal of Neuroscience. 2022;56(8):5137–5153. doi:10.1111/ejn.15812

109. Proulx CD, Hikosaka O, Malinow R. Reward processing by the lateral habenula in normal and depressive behaviors. Nat Neurosci. 2014;17(9):1146–1152. doi:10.1038/nn.3779

110. Majumdar A, Capetillo-Zarate E, Cruz D, Gouras GK, Maxfield FR. Degradation of Alzheimer’s amyloid fibrils by microglia requires delivery of ClC-7 to lysosomes. Mol Biol Cell. 2011;22(10):1664–1676. doi:10.1091/mbc.E10-09-0745

111. Wogram E, Wendt S, Matyash M, Pivneva T, Draguhn A, Kettenmann H. Satellite microglia show spontaneous electrical activity that is uncorrelated with activity of the attached neuron. Eur J Neurosci. 2016;43(11):1523–1534. doi:10.1111/ejn.13256

112. Pizzino G, Irrera N, Cucinotta M, et al. Oxidative Stress: Harms and Benefits for Human Health. Oxid Med Cell Longev. 2017;2017:8416763. doi:10.1155/2017/8416763

113. Maalouf M, Sullivan PG, Davis L, Kim DY, Rho JM. KETONES INHIBIT MITOCHONDRIAL PRODUCTION OF REACTIVE OXYGEN SPECIES PRODUCTION FOLLOWING GLUTAMATE EXCITOTOXICITY BY INCREASING NADH OXIDATION. Neuroscience. 2007;145(1):256–264. doi:10.1016/j.neuroscience.2006.11.065

114. Salim S. Oxidative Stress and Psychological Disorders. Curr Neuropharmacol. 2014;12(2):140–147. doi:10.2174/1570159X11666131120230309

115. Lehmann ML, Weigel TK, Poffenberger CN, Herkenham M. The Behavioral Sequelae of Social Defeat Require Microglia and Are Driven by Oxidative Stress in Mice. J Neurosci. 2019;39(28):5594–5605. doi:10.1523/JNEUROSCI.0184-19.2019

116. Katoh M, Wu B, Nguyen HB, et al. Polymorphic regulation of mitochondrial fission and fusion modifies phenotypes of microglia in neuroinflammation. Sci Rep. 2017;7:4942. doi:10.1038/s41598-017-05232-0

117. Peruzzotti-Jametti L, Willis CM, Hamel R, Krzak G, Pluchino S. Metabolic Control of Smoldering Neuroinflammation. Front Immunol. 2021;12:705920. doi:10.3389/fimmu.2021.705920

118. Skulachev VP. Mitochondrial filaments and clusters as intracellular power-transmitting cables. Trends in Biochemical Sciences. 2001;26(1):23–29. doi:10.1016/S0968-0004(00)01735-7

119. Karbowski M, Youle RJ. Dynamics of mitochondrial morphology in healthy cells and during apoptosis. Cell Death Differ. 2003;10(8):870–880. doi:10.1038/sj.cdd.4401260

120. Collins TJ, Berridge MJ, Lipp P, Bootman MD. Mitochondria are morphologically and functionally heterogeneous within cells. EMBO J. 2002;21(7):1616–1627. doi:10.1093/emboj/21.7.1616

121. Lounas A, Lebrun A, Laflamme I, et al. A 3D analysis revealed complexe mitochondria morphologies in porcine cumulus cells. Sci Rep. 2022;12:15403. doi:10.1038/s41598-022-19723-2

122. Loving BA, Bruce KD. Lipid and Lipoprotein Metabolism in Microglia. Front Physiol. 2020;11. doi:10.3389/fphys.2020.00393

123. Chausse B, Kakimoto PA, Caldeira-da-Silva CC, et al. Distinct metabolic patterns during microglial remodeling by oleate and palmitate. Biosci Rep. 2019;39(4):BSR20190072. doi:10.1042/BSR20190072

124. Mauerer R, Walczak Y, Langmann T. Comprehensive mRNA Profiling of Lipid-Related Genes in Microglia and Macrophages Using Taqman Arrays. In: Armstrong D, ed. Lipidomics: Volume 2: Methods and Protocols. Methods in Molecular Biology^TM^. Humana Press; 2010:187–201. doi:10.1007/978-1-60761-325-1_10

125. Nugent AA, Lin K, van Lengerich B, et al. TREM2 Regulates Microglial Cholesterol Metabolism upon Chronic Phagocytic Challenge. Neuron. 2020;105(5):837–854.e9. doi:10.1016/j.neuron.2019.12.007

126. Gulbins A, Schumacher F, Becker KA, et al. Antidepressants act by inducing autophagy controlled by sphingomyelin–ceramide. Mol Psychiatry. 2018;23(12):2324–2346. doi:10.1038/s41380-018-0090-9

127. Oliveira TG, Chan RB, Bravo FV, et al. The impact of chronic stress on the rat brain lipidome. Mol Psychiatry. 2016;21(1):80–88. doi:10.1038/mp.2015.14

128. Miranda AM, Oliveira TG. Lipids under stress – a lipidomic approach for the study of mood disorders. BioEssays. 2015;37(11):1226–1235. doi:10.1002/bies.201500070

129. Li KW, Ganz AB, Smit AB. Proteomics of neurodegenerative diseases: analysis of human post-mortem brain. Journal of Neurochemistry. 2019;151(4):435–445. doi:10.1111/jnc.14603

130. Ma Y, Poole K, Goyette J, Gaus K. Introducing Membrane Charge and Membrane Potential to T Cell Signaling. Frontiers in Immunology. 2017;8. Accessed March 21, 2023. https://www.frontiersin.org/articles/10.3389/fimmu.2017.01513

131. Bruzzese A, Gil C, Dalton JAR, Giraldo J. Structural insights into positive and negative allosteric regulation of a G protein-coupled receptor through protein-lipid interactions. Sci Rep. 2018;8:4456. doi:10.1038/s41598-018-22735-6

132. Olofsson KE, Andersson L, Nilsson J, Björkbacka H. Nanomolar concentrations of lysophosphatidylcholine recruit monocytes and induce pro-inflammatory cytokine production in macrophages. Biochemical and Biophysical Research Communications. 2008;370(2):348–352. doi:10.1016/j.bbrc.2008.03.087

133. Yang LV, Radu CG, Wang L, Riedinger M, Witte ON. Gi-independent macrophage chemotaxis to lysophosphatidylcholine via the immunoregulatory GPCR G2A. Blood. 2005;105(3):1127–1134. doi:10.1182/blood-2004-05-1916

134. Chen S, Wei C, Gao P, et al. Effect of Allium macrostemon on a rat model of depression studied by using plasma lipid and acylcarnitine profiles from liquid chromatography/mass spectrometry. Journal of Pharmaceutical and Biomedical Analysis. 2014;89:122–129. doi:10.1016/j.jpba.2013.10.045

135. Weng R, Shen S, Burton C, et al. Lipidomic profiling of tryptophan hydroxylase 2 knockout mice reveals novel lipid biomarkers associated with serotonin deficiency. Anal Bioanal Chem. 2016;408(11):2963–2973. doi:10.1007/s00216-015-9256-3

136. Lee LHW, Shui G, Farooqui AA, Wenk MR, Tan CH, Ong WY. Lipidomic analyses of the mouse brain after antidepressant treatment: evidence for endogenous release of long-chain fatty acids? International Journal of Neuropsychopharmacology. 2009;12(7):953–964. doi:10.1017/S146114570900995X

137. Neher JJ, Neniskyte U, Zhao JW, Bal-Price A, Tolkovsky AM, Brown GC. Inhibition of Microglial Phagocytosis Is Sufficient To Prevent Inflammatory Neuronal Death. The Journal of Immunology. 2011;186(8):4973–4983. doi:10.4049/jimmunol.1003600

138. Kurematsu C, Sawada M, Ohmuraya M, et al. Synaptic pruning of murine adult-born neurons by microglia depends on phosphatidylserine. J Exp Med. 2022;219(4):e20202304. doi:10.1084/jem.20202304

139. Scott-Hewitt N, Perrucci F, Morini R, et al. Local externalization of phosphatidylserine mediates developmental synaptic pruning by microglia. The EMBO Journal. 2020;n/a(n/a):e105380. doi:10.15252/embj.2020105380

140. Yin W, Gallagher NR, Sawicki CM, McKim DB, Godbout JP, Sheridan JF. Repeated Social Defeat in Female Mice Induces Anxiety-Like Behavior Associated with Enhanced Myelopoiesis and Increased Monocyte Accumulation in the Brain. Brain Behav Immun. 2019;78:131–142. doi:10.1016/j.bbi.2019.01.015

141. van Doeselaar L, Yang H, Bordes J, et al. Chronic social defeat stress in female mice leads to sex-specific behavioral and neuroendocrine effects. Stress. 2021;24(2):168–180. doi:10.1080/10253890.2020.1864319

142. Kinsey SG, Bailey MT, Sheridan JF, Padgett DA. The Inflammatory Response to Social Defeat is Increased in Older Mice. Physiol Behav. 2008;93(3):628–636. doi:10.1016/j.physbeh.2007.11.003

143. Kumar NK, Merrill JD, Carlson S, German J, Yancy WS. Adherence to Low-Carbohydrate Diets in Patients with Diabetes: A Narrative Review. Diabetes Metab Syndr Obes. 2022;15:477–498. doi:10.2147/DMSO.S292742

144. Fortier M, Castellano C, StLPierre V, et al. A ketogenic drink improves cognition in mild cognitive impairment: Results of a 6Lmonth RCT. Alzheimers Dement. 2021;17(3):543–552. doi:10.1002/alz.12206

145. Croteau E, Castellano CA, Richard MA, et al. Ketogenic Medium Chain Triglycerides Increase Brain Energy Metabolism in Alzheimer’s Disease. Journal of Alzheimer’s Disease. 2018;64(2):551–561. doi:10.3233/JAD-180202

