## Supplementary Material for "Ketogenic diet alters microglial morphology and changes the hippocampal lipidomic profile distinctively in stress susceptible *versus* resistant male mice upon repeated social defeat"

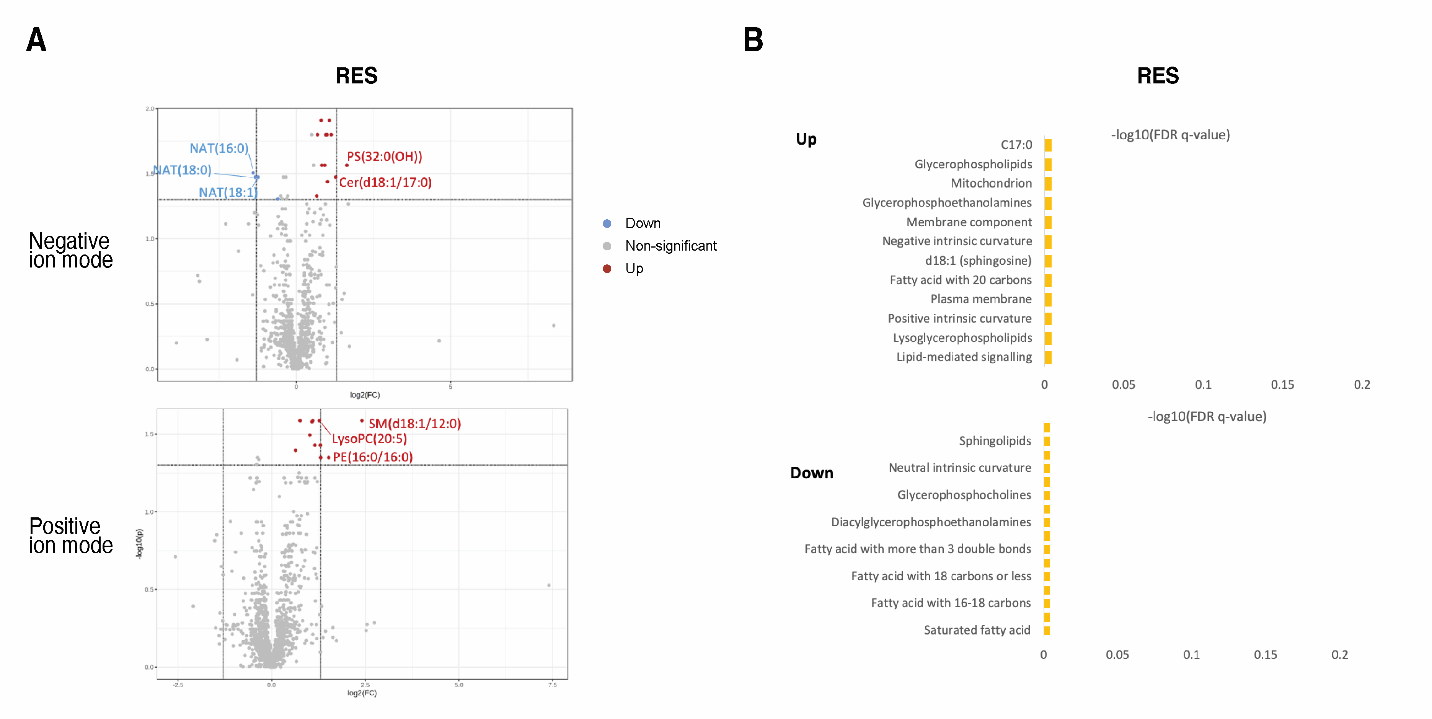

**Figure S1: Effect of KD treatment on hippocampal lipidomic profiles in stress-resistant mice. A)** Volcano plots illustrating differentially upregulated (red) and downregulated (blue) lipids in whole hippocampus of stress-resistant (RES) mice fed a ketogenic diet (KD), relative to a control diet (CD) (n = 4 mice per group). Positive and negative ion mode datasets were analyzed separately using two-sample t-tests, with a fold change of >1.5 and an FDR-adjusted p-value (q) ≤ 0.05 that were considered significant. Cer, ceramide; NAT, N-acyltaurine; PC, phosphatidylcholine; PE, phosphatdiylethanolamine; PS, phosphatidylserine; SM, sphingomyelin. B) Ontological analysis of differentially regulated lipids between KD and CD diets in RES mice. Differentially regulated lipids were annotated by matching to the human metabolome database (HMDB), followed by analysis in LION software in ranking mode. Lipid ontology terms with a fold change of >1.3 in either direction q ≤ 0.05 were considered significantly enriched (top) or underrepresented (bottom) in response to a ketogenic diet. No significantly enriched or underrepresented terms were identified in RES mice.

**Table S1: Hippocampal lipids differentially regulated by a ketogenic diet in non-stressed and stress-susceptible mice.** Differentially regulated hippocampal lipids between non-stressed control (CTRL), stress-susceptible (SUS) and stress-resistant (RES) mice fed either a ketogenic (KD) or control diet (CD) prior to repeated social defeat stress exposure (n = 4 mice per treatment). Positive and negative-ion mode lipidomic datasets were analyzed separately using two-sample t-tests, with features considered significant with a fold change >1.5 in either direction and a false-discovery rate (FDR)-adjusted p-value ≤0.05. Features identified as significant were annotated by matching m/z ratios and retention times to the human metabolome database (HMDB). Displayed here are lipids successfully matched to an identifier. Cer, ceramide; CL, cardiolipin; HexCer, hexosylceramide; FA, fatty acid; GlcCer, glucosylceramide; NAT, N-acyltaurine; PA, phosphatidic acid; PC, phosphatidylcholine; PE, phosphatidylethanolamine; PG, phosphatidylglycerol; PI, phosphatidylinositol; PIP, phosphatidylinositol-monophosphate; PG, phosphatidylglycerol; PS, phosphatidylserine; SHexCer, sulfated hexosylceramide; SM, sphingomyelin; SQDG, sulfoquinovosyl-diacylglycerol.

| m/z | **Retention Time (s)** | **Lipid Species** | **Formula** | **Fold Change** | log2(Fold Change) | FDR-adjusted p-value (q) | -log10  (q) |
| --- | --- | --- | --- | --- | --- | --- | --- |
| Negative Ion-Mode, KD-CTRL vs CD-CTRL: Up |  |  |  |  |  |  |  |
| 731.48458 | 12.25 | PA(P-16:0/PGF1alpha) | C39H72O10P | 4.1255 | 2.0446 | 0.039179 | 1.4069 |
| 719.53212 | 9.73 | SM(d18:1/14:0) | C37H75N2O6P | 2.5655 | 1.3593 | 0.032786 | 1.4843 |
| 736.51133 | 10.16 | PS(O-32:0(OH)) | C38H75NO10P | 2.4277 | 1.2796 | 0.024737 | 1.6067 |
| 554.32371 | 6.95 | LysoPE(22:2) | C29H49NO7P | 2.043 | 1.0307 | 0.020664 | 1.6848 |
| 826.50971 | 9.86 | PE(18:3/TXB2) | C43H74NO12P | 1.9026 | 0.92798 | 0.020664 | 1.6848 |
| 427.32055 | 8.54 | 1,24(S)-Dihydroxyvitamin D2 | C28H43O3 | 1.8388 | 0.87876 | 0.039409 | 1.4044 |
| 427.32055 | 8.54 | Ercalcitriol | C28H43O3 | 1.8388 | 0.87876 | 0.039409 | 1.4044 |
| 494.32399 | 6.95 | LysoPE(19:0) | C24H49NO7P | 1.8112 | 0.85691 | 0.032786 | 1.4843 |
| 528.30821 | 6.34 | LysoPE(22:4) | C27H47NO7P | 1.6458 | 0.71878 | 0.036028 | 1.4434 |
| 664.51113 | 11.66 | Cer(d18:1/22:4)-OH | C40H71NO4 | 1.6397 | 0.71346 | 0.039409 | 1.4044 |
| 764.5424 | 10.78 | PS(O-18:0/16:0)-OH | C40H79NO10P | 1.6237 | 0.69932 | 0.032786 | 1.4843 |
| 466.32873 | 7.57 | PE(O-18:0) | C23H49NO6P | 1.5409 | 0.62375 | 0.032786 | 1.4843 |
| 509.3428 | 7.27 | LysoPE(20:0) | C25H52NO7P | 1.539 | 0.622 | 0.02916 | 1.5352 |
| Positive Ion-Mode, KD-CTRL vs CD-CTRL: Up |  |  |  |  |  |  |  |
| 647.51298 | 8.93 | SM(d18:1/12:0) | C35H72N2O6P | 6.2906 | 2.6532 | 0.002985 | 2.5251 |
| 673.52879 | 9.04 | SM(d18:1/14:1) | C37H74N2O6P | 3.4151 | 1.7719 | 0.016368 | 1.786 |
| 532.33944 | 7.48 | LysoPE(20:0) | C25H52NO7PNa | 2.9601 | 1.5656 | 0.037013 | 1.4316 |
| 510.35558 | 6.95 | LysoPA(22:1) | C25H53NO7P | 2.1713 | 1.1186 | 0.020655 | 1.685 |
| 510.35558 | 6.95 | LysoPC(17:0/0:0) | C25H53NO7P | 2.1713 | 1.1186 | 0.020655 | 1.685 |
| 675.54431 | 9.6 | SM(d18:1/14:0) | C37H76N2O6P | 1.967 | 0.97603 | 0.023756 | 1.6242 |
| 566.32214 | 6.35 | LysoPC(0:0/20:4(5Z,8Z,11Z,14Z)) | C28H50NO7PNa | 1.9383 | 0.95481 | 0.023756 | 1.6242 |
| 722.55591 | 10.8 | GlcCer(d18:1/16:0) | C40H77NO8Na | 1.7737 | 0.82675 | 0.047841 | 1.3202 |
| 506.36079 | 7.27 | LysoPC(P-18:1) | C26H53NO6P | 1.6955 | 0.76169 | 0.020655 | 1.685 |
| 478.32981 | 6.64 | LysoPC(16:2) | C24H49NO6P | 1.6524 | 0.72456 | 0.023756 | 1.6242 |
| Negative Ion-Mode, KD-CTRL vs CD-CTRL: Down |  |  |  |  |  |  |  |
| 571.28676 | 6.78 | LysoPI(16:0/0:0) | C25H48O12P | 0.20279 | -2.302 | 0.048174 | 1.3172 |
| 597.30257 | 6.94 | LysoPI(18:1(9Z)/0:0) | C27H50O12P | 0.23258 | -2.1042 | 0.033298 | 1.4776 |
| 619.28695 | 6.49 | PI(20:4(5Z,8Z,11Z,14Z)/0:0) | C29H48O12P | 0.27653 | -1.8545 | 0.032786 | 1.4843 |
| 590.24796 | 6.49 | LysoPS(22:6) | C28H43NO9P | 0.28252 | -1.8236 | 0.04452 | 1.3514 |
| 802.54765 | 12.18 | PS(18:2/19:0) | C43H80NO10P | 0.39765 | -1.3304 | 0.039179 | 1.4069 |
| 509.28714 | 6.98 | LysoPG(18:1(9Z)/0:0) | C24H46O9P | 0.43901 | -1.1877 | 0.049019 | 1.3096 |
| 568.26634 | 6.49 | LysoPS(22:6) | C28H43NO9P | 0.47108 | -1.086 | 0.032786 | 1.4843 |
| 916.52845 | 12.29 | PS(22:6-2OH/20:0) | C48H82NO12P | 0.47425 | -1.0763 | 0.020664 | 1.6848 |
| 912.51586 | 10.66 | PC(20:4/PGD2) | C48H80NO11P | 0.48896 | -1.0322 | 0.032786 | 1.4843 |
| 829.64964 | 13.94 | PE(24:0/18:1) | C47H92NO8P | 0.65703 | -0.60598 | 0.032786 | 1.4843 |
| Positive Ion-Mode, KD-CTRL vs CD-CTRL: Down |  |  |  |  |  |  |  |
| n/a |  |  |  |  |  |  |  |
| Negative Ion-Mode, KD-SUS vs CD-SUS: Up |  |  |  |  |  |  |  |
| 824.54494 | 11.11 | PE(18:1-2OH/22:4) | C45H79NO10P | 4.3713 | 2.1281 | 0.033401 | 1.4762 |
| 736.51133 | 10.16 | PS(16:0/16:0-OH) | C38H75NO10P | 2.6217 | 1.3905 | 0.033401 | 1.4762 |
| 550.51852 | 11.66 | Cer(d18:1/17:0) | C35H68NO3 | 2.3755 | 1.2482 | 0.037271 | 1.4286 |
| 427.32055 | 8.54 | 1,24(S)-Dihydroxyvitamin D2 | C28H43O3 | 2.2759 | 1.1865 | 0.048622 | 1.3132 |
| 427.32055 | 8.54 | Ercalcitriol | C28H43O3 | 2.2759 | 1.1865 | 0.048622 | 1.3132 |
| 837.53491 | 11.39 | PS(18:1/22:5) | C46H78O11P | 2.242 | 1.1648 | 0.046847 | 1.3293 |
| 830.49451 | 10.66 | PS(18:2/22:6) | C46H73NO10P | 1.9309 | 0.94925 | 0.017065 | 1.7679 |
| 554.32371 | 6.95 | LysoPE(22:2) | C29H49NO7P | 1.8057 | 0.85253 | 0.033401 | 1.4762 |
| 1035.66007 | 6.64 | LysoPC(16:0/0:0) | C57H96O14P | 1.774 | 0.82703 | 0.043826 | 1.3583 |
| 704.52104 | 10.8 | PE(18:0/15:0) | C38H75NO8P | 1.6942 | 0.76058 | 0.033401 | 1.4762 |
| 631.47056 | 7.47 | PA(18:1/14:0) | C35H68O7P | 1.6679 | 0.73799 | 0.043826 | 1.3583 |
| 764.5424 | 10.78 | PS(16:0/18:0-OH) | C40H79NO10P | 1.6649 | 0.73547 | 0.033401 | 1.4762 |
| 748.49024 | 10.65 | PE(15:0/20:3) | C38H70NO13 | 1.6615 | 0.7325 | 0.043826 | 1.3583 |
| 783.49797 | 10.97 | PI(14:0/16:0-OH) | C39H76O13P | 1.648 | 0.72073 | 0.033401 | 1.4762 |
| 806.49534 | 10.9 | PS(16:0/22:6) | C44H73NO10P | 1.5446 | 0.62723 | 0.044852 | 1.3482 |
| 279.23244 | 7.51 | Linoleic acid | C18H31O2 | 1.5128 | 0.59719 | 0.043826 | 1.3583 |
| Positive Ion-Mode, KD-SUS vs CD-SUS: Up |  |  |  |  |  |  |  |
| 722.55591 | 10.8 | GlcCer(d18:1/16:0) | C40H77NO8Na | 1.8948 | 0.92204 | 0.0092407 | 2.0343 |
| 717.59204 | 10.63 | SM(d18:1/17:0) | C40H82N2O6P | 1.7028 | 0.76788 | 0.014809 | 1.8295 |
| 1082.85331 | 15.18 | PC(58:8) | C66H117NO8P | 1.6732 | 0.74263 | 0.02905 | 1.5369 |
| 1540.12488 | 10.57 | CL(16:0/18:0/18:2/24:0) | C87H161O17P2 | 1.6456 | 0.71861 | 0.030935 | 1.5096 |
| Negative Ion-Mode, KD-SUS vs CD SUS: Down |  |  |  |  |  |  |  |
| 571.28676 | 6.78 | LysoPI(16:0/0:0) | C25H48O12P | 0.10399 | -3.2654 | 0.045107 | 1.3458 |
| 643.28658 | 6.49 | LysoPI(22:6) | C31H48O12P | 0.20463 | -2.2889 | 0.0122 | 1.9136 |
| 362.2361 | 6.92 | NAT(16:0) | C18H36NO4S | 0.28227 | -1.8249 | 0.037271 | 1.4286 |
| 555.27032 | 6.63 | LysoPG(22:6) | C28H44O9P | 0.28751 | -1.7983 | 0.036452 | 1.4383 |
| 509.28714 | 6.98 | LysoPG(18:1(9Z)/0:0) | C24H46O9P | 0.30075 | -1.7334 | 0.014124 | 1.85 |
| 544.26595 | 6.5 | LysoPS(20:4) | C26H43NO9P | 0.31637 | -1.6603 | 0.048622 | 1.3132 |
| 522.28256 | 6.85 | LysoPS(18:1(9Z)/0:0) | C24H45NO9P | 0.32018 | -1.643 | 0.020234 | 1.6939 |
| 483.27148 | 6.94 | LysoPG(16:0/0:0) | C22H44O9P | 0.32697 | -1.6128 | 0.01649 | 1.7828 |
| 840.60962 | 13.8 | PE(18:1-O/24:1) | C47H87NO9P | 0.42541 | -1.2331 | 0.043826 | 1.3583 |
| 854.54997 | 13.23 | PS(40:4(OH)) | C46H81NO11P | 0.46798 | -1.0955 | 0.033401 | 1.4762 |
| 536.29778 | 7.31 | LysoPS(19:1/0:0) | C25H47NO9P | 0.48894 | -1.0323 | 0.0122 | 1.9136 |
| 510.28224 | 7.25 | LysoPS(17:0/0:0) | C23H45NO9P | 0.51162 | -0.96685 | 0.043826 | 1.3583 |
| 808.53761 | 11.96 | SM(d18:1/20:4-2OH) | C41H78NO14 | 0.55684 | -0.84468 | 0.0122 | 1.9136 |
| 772.54716 | 12.81 | PS(18:0/18:2-OH) | C42H79NO9P | 0.58624 | -0.77043 | 0.033401 | 1.4762 |
| 688.48984 | 10.75 | PE(18:1(9Z)/14:0) | C37H71NO8P | 0.63406 | -0.6573 | 0.0122 | 1.9136 |
| Positive Ion-Mode, KD-SUS vs CD-SUS: Down |  |  |  |  |  |  |  |
| 592.26479 | 6.5 | LysoPS(22:6) | C28H44NO9PNa | 0.26199 | -1.9324 | 0.02905 | 1.5369 |
| 915.71688 | 13.19 | PE(46:3(OH)) | C51H100N2O9P | 0.38813 | -1.3654 | 0.0092407 | 2.0343 |
| 831.62368 | 13.18 | PE(18:2(O)/22:0) | C45H88N2O9P | 0.4031 | -1.3108 | 0.049882 | 1.3021 |
| 903.71716 | 13.78 | PC(24:0/18:1(9Z)-O) | C50H100N2O9P | 0.40777 | -1.2942 | 0.02905 | 1.5369 |
| 917.73289 | 13.78 | PE(46:2(OH)) | C51H102N2O9P | 0.42098 | -1.2482 | 0.014809 | 1.8295 |
| 805.60789 | 13.22 | PC(18:1-O/17:0) | C43H86N2O9P | 0.43625 | -1.1968 | 0.02905 | 1.5369 |
| 786.56571 | 12.58 | PA(40:4(OH)) | C43H81NO9P | 0.43991 | -1.1847 | 0.049061 | 1.3093 |
| 890.63993 | 13.22 | SHexCer(d18:1/24:1) | C48H92NO11S | 0.44966 | -1.1531 | 0.0058841 | 2.2303 |
| 816.61222 | 13.78 | PE(18:1-O/22:0) | C45H87NO9P | 0.45138 | -1.1476 | 0.02905 | 1.5369 |
| 838.59471 | 13.77 | PS(16:0/22:2) | C45H86NO9PNa | 0.45161 | -1.1468 | 0.02905 | 1.5369 |
| 875.68589 | 13.22 | PC(18:1-O/22:0) | C48H96N2O9P | 0.45754 | -1.128 | 0.010664 | 1.9721 |
| 538.31419 | 13.22 | LysoPS(19:1) | C25H49NO9P | 0.45886 | -1.1239 | 0.02905 | 1.5369 |
| 889.7015 | 13.22 | PE(44:2(OH)) | C49H98N2O9P | 0.46929 | -1.0914 | 0.014809 | 1.8295 |
| 690.50759 | 10.76 | PE(18:1/14:0) | C37H73NO8P | 0.60632 | -0.72185 | 0.02905 | 1.5369 |
| 690.50759 | 10.76 | PA(16:0/18:2) | C37H73NO8P | 0.60632 | -0.72185 | 0.02905 | 1.5369 |
| 770.51031 | 10.64 | SHexCer(t16:0/16:0) | C38H76NO12S | 0.62595 | -0.67588 | 0.033263 | 1.478 |
| Negative Ion-Mode, KD-RES vs ND-RES: Up |  |  |  |  |  |  |  |
| 736.51133 | 10.16 | PS(O-32:0(OH)) | C38H75NO10P | 3.0977 | 1.6312 | 0.027207 | 1.5653 |
| 550.51852 | 11.66 | Cer(d18:1/17:0) | C35H68NO3 | 2.4159 | 1.2726 | 0.033595 | 1.4737 |
| 664.51113 | 11.66 | Cer(d20:1/20:4-OH(16R)) | C40H70NO4 | 2.1856 | 1.128 | 0.015865 | 1.7996 |
| 466.29258 | 7.01 | PE(17:0/0:0) | C22H45NO7P | 2.0061 | 1.0044 | 0.036487 | 1.4379 |
| 764.5424 | 10.78 | PS(O-34:0(OH)) | C40H79NO10P | 1.9842 | 0.98854 | 0.015865 | 1.7996 |
| 596.5238 | 11.66 | Cer(d18:1/18:0) | C36H71NO3 | 1.9376 | 0.95428 | 0.015892 | 1.7988 |
| 749.49359 | 10.66 | PE(22:6/15:0) | C42H72NO8P | 1.8893 | 0.91787 | 0.027207 | 1.5653 |
| 872.53426 | 11.18 | SHexCer(t18:1/19:0) | C43H83NO10S | 1.7424 | 0.8011 | 0.012317 | 1.9095 |
| 622.46407 | 10.56 | NAT(32:0) | C24H49NO4S | 1.6085 | 0.68576 | 0.015865 | 1.7996 |
| 753.52458 | 11.47 | PC(16:0/18:4) | C42H76NO8P | 1.5803 | 0.66023 | 0.046917 | 1.3287 |
| Positive Ion-Mode, KD-RES vs ND-RES: Up |  |  |  |  |  |  |  |
| 647.51298 | 8.93 | SM(d18:1/12:0) | C35H72N2O6P | 5.3029 | 2.4068 | 0.0259 | 1.5867 |
| 692.52344 | 10.11 | PE(16:0/16:0) | C37H75NO8P | 2.8641 | 1.5181 | 0.044687 | 1.3498 |
| 542.32192 | 6.39 | LysoPC(20:5) | C28H49NO7P | 2.4141 | 1.2715 | 0.0259 | 1.5867 |
| 548.3713 | 6.94 | LysoPC(20:2) | C28H55NO7P | 2.1355 | 1.0946 | 0.0259 | 1.5867 |
| 722.55591 | 10.8 | GlcCer(d18:1/16:0) | C40H77NO8Na | 2.0243 | 1.0174 | 0.032105 | 1.4934 |
| 647.51298 | 8.93 | SM(d18:1/17:0) | C41H84O7P | 1.6982 | 0.76405 | 0.0259 | 1.5867 |
| 692.52344 | 10.11 | PE(18:0/22:5) | C45H81NO8P | 1.6835 | 0.75148 | 0.0259 | 1.5867 |
| Negative Ion-Mode, KD-RES vs ND-RES: Down |  |  |  |  |  |  |  |
| 362.2361 | 6.92 | NAT(16:0) | C18H36NO4S | 0.37841 | -1.402 | 0.031181 | 1.5061 |
| 390.26722 | 7.63 | NAT(18:0) | C20H40NO4S | 0.39799 | -1.3292 | 0.033595 | 1.4737 |
| 388.25157 | 7.09 | NAT(18:1) | C20H38NO4S | 0.42252 | -1.2429 | 0.033595 | 1.4737 |
| 813.52424 | 10.84 | PG(18:0/20:4)-OH(20)) | C44H78O11P | 0.65936 | -0.60085 | 0.049391 | 1.3064 |

**Table S2: Lipid ontology term enrichment analyses.** Ontological analyses of differentially regulated lipids between (CTRL), stress-susceptible (SUS) and stress-resistant (RES) mice fed a ketogenic (KD) compared to control diet (CD), and exposed to repeated social defeat stress (n = 4 mice per treatment). Lipids identified as differentially regulated by two-sample t-test analyses were annotated by matching to the human metabolome database (HMDB), and annotated lipids were analyzed using LION software (v.2020.07.14) in ranking mode. All matched lipid ontology terms are displayed below. Terms with a fold change of >1.3 in either direction with a false-discovery rate (FDR)-adjusted p-value (q) of ≤0.05 were considered significantly enriched or underrepresented. No significantly enriched or underrepresented groups of lipids were identified in stress-resilient mice.

| Term ID | **Description** | **Annotated** | **p-value** | **FDR q-value** | **Effect Size** | **Regulated** |
| --- | --- | --- | --- | --- | --- | --- |
| **KD-CTRL vs ND-CTRL** |  |  |  |  |  |  |
| **LION:0000093** | headgroup with negative charge | 10 | 6.10E-05 | 0.00244 | -0.8 | DOWN |
| LION:0000046 | monoacylglycerophosphoinositols [GP0605] | 3 | 0.00055 | 0.011 | -1 | DOWN |
| LION:0000095 | headgroup with positive charge / zwitter-ion | 16 | 0.00396 | 0.04712 | 0.615384615 | UP |
| LION:0000084 | ceramide phosphocholines (sphingomyelins) [SP0301] | 3 | 0.00547 | 0.04712 | 0.923076923 | UP |
| LION:0000013 | glycerophosphoserines [GP03] | 4 | 0.00589 | 0.04712 | -0.84 | DOWN |
| LION:0002937 | C22:6 | 3 | 0.03065 | 0.20433 | -0.807692308 | DOWN |
| LION:0080947 | d18:1 (sphingosine) | 5 | 0.0514 | 0.29371 | 0.625 | UP |
| LION:0002976 | fatty acid with more than 3 double bonds | 9 | 0.06007 | 0.30035 | -0.5 | DOWN |
| LION:0002957 | fatty acid with 18 carbons | 8 | 0.08204 | 0.36462 | -0.494047619 | DOWN |
| LION:0002967 | polyunsaturated fatty acid | 11 | 0.15244 | 0.52067 | -0.409090909 | DOWN |
| LION:0000465 | neutral intrinsic curvature | 4 | 0.1562 | 0.52067 | -0.56 | DOWN |
| LION:0002922 | C18:1 | 4 | 0.1562 | 0.52067 | -0.56 | DOWN |
| LION:0000003 | glycerophospholipids [GP] | 23 | 0.1748 | 0.53785 | -0.47826087 | DOWN |
| LION:0000011 | glycerophosphoethanolamines [GP02] | 7 | 0.24233 | 0.56891 | 0.409090909 | UP |
| LION:0002966 | fatty acid with less than 2 double bonds | 19 | 0.26203 | 0.56891 | 0.373684211 | UP |
| LION:0002977 | fatty acid with 3-5 double bonds | 6 | 0.26421 | 0.56891 | -0.434782609 | DOWN |
| LION:0002972 | fatty acid with 4 double bonds | 5 | 0.27658 | 0.56891 | -0.458333333 | DOWN |
| LION:0001737 | average transition temperature | 3 | 0.28134 | 0.56891 | 0.551282051 | UP |
| LION:0000466 | positive intrinsic curvature | 15 | 0.29698 | 0.56891 | -0.328571429 | DOWN |
| LION:0000599 | lysoglycerophospholipids | 15 | 0.29698 | 0.56891 | -0.328571429 | DOWN |
| LION:0000042 | monoacylglycerophosphoethanolamines [GP0205] | 4 | 0.29868 | 0.56891 | 0.48 | UP |
| LION:0002945 | fatty acid with more than 18 carbons | 14 | 0.44946 | 0.6768 | -0.3 | DOWN |
| LION:0002968 | saturated fatty acid | 14 | 0.44946 | 0.6768 | 0.3 | UP |
| LION:0012080 | endoplasmic reticulum (ER) | 17 | 0.45214 | 0.6768 | -0.299019608 | DOWN |
| LION:0002948 | fatty acid with 16-18 carbons | 13 | 0.45691 | 0.6768 | -0.298076923 | DOWN |
| LION:0002949 | fatty acid with 19-21 carbons | 7 | 0.46377 | 0.6768 | -0.344155844 | DOWN |
| LION:0002929 | C20:4 | 3 | 0.50739 | 0.6768 | -0.461538462 | DOWN |
| LION:0000464 | negative intrinsic curvature | 5 | 0.50843 | 0.6768 | 0.375 | UP |
| LION:0000010 | glycerophosphocholines [GP01] | 6 | 0.51561 | 0.6768 | 0.347826087 | UP |
| LION:0000034 | monoacylglycerophosphocholines [GP0105] | 4 | 0.52166 | 0.6768 | 0.4 | UP |
| LION:0012082 | plasma membrane | 8 | 0.52452 | 0.6768 | -0.30952381 | DOWN |
| LION:0002959 | fatty acid with 20 carbons | 5 | 0.57514 | 0.7081 | -0.35 | DOWN |
| LION:0012009 | lipid-mediated signalling | 17 | 0.58418 | 0.7081 | -0.269607843 | DOWN |
| LION:0012081 | mitochondrion | 8 | 0.74389 | 0.87516 | 0.255952381 | UP |
| LION:0002950 | fatty acid with 22-24 carbons | 8 | 0.79296 | 0.90624 | -0.244047619 | DOWN |
| LION:0012010 | membrane component | 27 | 0.84236 | 0.93596 | -0.407407407 | DOWN |
| LION:0002961 | fatty acid with 22 carbons | 7 | 0.87893 | 0.95019 | -0.227272727 | DOWN |
| LION:0000100 | fatty acid with 18 carbons or less | 16 | 0.90399 | 0.95157 | 0.1875 | UP |
| LION:0002882 | C16:0 | 4 | 0.94543 | 0.96842 | -0.25 | DOWN |
| LION:0002969 | monounsaturated fatty acid | 6 | 0.96842 | 0.96842 | -0.195652174 | DOWN |
